# The circadian clock controls temporal and spatial patterns of floral development in sunflower

**DOI:** 10.1101/2022.06.30.498369

**Authors:** Carine M. Marshall, Veronica L. Thompson, Nicky M. Creux, Stacey L. Harmer

## Abstract

Biological rhythms are ubiquitous. They can be generated by circadian oscillators, which produce daily rhythms in physiology and behavior, as well as by developmental oscillators such as the segmentation clock, which produces modular developmental units in a periodic fashion. Here, we show that the circadian clock controls the timing of late-stage floret development, or anthesis, in domesticated sunflower. In these plants, what appears to be a single inflorescence consists of up to thousands of individual florets tightly packed onto a capitulum disk. While early floret development occurs continuously across capitula to generate iconic spiral phyllotaxy, during anthesis floret development occurs in discrete ring-like pseudowhorls with up to hundreds of florets undergoing simultaneous maturation. We demonstrate circadian regulation of floral organ growth and show that the effects of light on this process are time-of-day dependent. Disruption of circadian rhythms in floral organ development causes loss of pseudowhorl formation. Thus, we show that the sunflower circadian clock acts in concert with environmental response pathways to tightly synchronize the anthesis of hundreds of florets each day, generating spatial patterns on the developing capitulum disk. This coordinated mass release of floral rewards at predictable times of day likely promotes pollinator visits and plant reproductive success.

## Introduction

From the earliest stages of cellular differentiation to reproductive maturity, spatio-temporal regulation of developmental transitions plays an important role in biological success. In some cases, internal temporal rhythms are used to spatially organize a body plan. For example, in vertebrate embryos a molecular ‘segmentation clock’ acts at regular intervals to divide the developing body axis into discrete blocks of cells that later differentiate into individual somites (Richmond and Oates, 2012). In plants, the regular spacing of lateral roots along the length of the primary root axis is controlled by a ‘root clock’ that generates a temporally oscillating developmental signal (Xuan et al., 2020). Thus in both plants and animals, internal body clocks can generate serially-repeated organs. These types of clocks are distinct from the circadian oscillator, the biological clock found in most eukaryotes and some prokaryotes (Mosier and Hurley, 2021). Circadian clocks coordinate development on daily and seasonal time scales. In both plants and animals, the circadian clock can integrate environmental cues such as day length and temperature to time the transition to reproductive stages to the appropriate season (Ikegami and Yoshimura, 2012; Inoue et al., 2018). On a diel scale, the circadian clock can coordinate developmental processes with environmental transitions associated with the earth’s rotation, such as leaf movement in plants, eclosion in *Drosophila*, and conidiation in *Neurospora* (Brady et al., 2021; Larrondo and Canessa, 2018; McClung, 2019). However, a role for the circadian clock in the spatial patterning of development has not previously been reported.

Circadian clocks in eukaryotes are cell-autonomous systems made up of transcriptional-translational feedback loops that create rhythms of gene expression to regulate biological timing. Circadian rhythms, biological processes regulated by the circadian clock, meet several fundamental criteria (Rensing and Ruoff, 2002). First, they are stably entrained to environmental cues. Second, they persist in free running environmental conditions. Third, circadian rhythms are temperature compensated such that they maintain a uniform period over a range of ambient temperatures, in contrast to the general rule that the rate of enzymatic processes changes with temperature. In addition, the circadian clock gates certain biological processes so that responsiveness to environmental cues is time-of-day dependent. For example, studies in humans show the timing of drug treatment can be critical to creating an effective therapeutic response (Ruan et al., 2021). A functioning circadian clock whose biological rhythms match that of the environment provides a fitness advantage for many organisms, including plants (Dodd et al., 2005; Michael et al., 2003), animals (Miller et al., 2004; Emerson et al., 2008; Spoelstra et al., 2016), bacteria (Ouyang et al., 1998), and fungi (Koritala et al., 2020).

One important way the circadian clock promotes fitness is by controlling timing of reproductive development, for example by allowing organisms to distinguish between the long and short days of different seasons. In plants, the circadian system mediates the effects of day length during the transition from vegetative to reproductive growth, usually referred to as flowering time (Imaizumi and Kay, 2006). The circadian clock also regulates daily aspects of floral development, for example coordinating daily rhythms in scent emission and floral opening and closing with pollinator activity (Yon et al., 2017; Inoue et al., 2018; Muroya et al., 2021). In addition, the circadian clock is implicated in the uniform eastward orientation observed in mature sunflower plants; this eastward orientation was shown to enhance both male and female reproductive fitness (Atamian et al., 2016; Creux et al., 2021).

Domesticated sunflower is an excellent model system to study the temporal regulation of floral development. Sunflowers are members of the Asteraceae, a highly successful family comprised of about 30,000 distinct species that are widely dispersed around the world (Funk et al., 2009). Asteraceae are characterized by the arrangement of individual florets together in a complex, compressed inflorescence called the capitulum. Individual florets are arranged on the capitulum disk in spiral patterns, a classic type of phyllotaxis observed in many plants (Figures 1A, 1B). These phyllotactic spirals, called parastichies, are generated by the precise specification of floral primordia, first at the outer rim and then moving into the center of the capitulum disk (Zhang et al., 2021). In domesticated sunflower, specification of the hundreds or even thousands of floret primordia found on a single capitulum takes approximately 10-14 days (Marc and Palmer, 1981; Palmer and Steer, 1985), followed approximately 20 days later by the onset of anthesis, or floral opening, of the first florets (Lindström and Hernández, 2015). Anthesis of all florets on a capitulum head typically occurs over 5-10 days (Seiler, 2015). Intriguingly, the timing of anthesis does not follow the gradual and continuous pattern of development seen during specification of floret primordia on the developing capitulum (Marc and Palmer, 1981; Zhang et al., 2021). Instead, discrete developmental boundaries are observed with obvious rings of florets at distinct stages of anthesis (Figures 1E, 1F). Each ring, or pseudowhorl, undergoes anthesis on successive days. While previous studies examined the effects of hormones and environmental factors on the timing of development of florets within a pseudowhorl (Baroncelli et al., 1990; Lobello et al., 2000), the mechanisms governing the conversion of the continuous spiral pattern of development seen early in floral development (Figure 1A) to the ringlike patterns seen during anthesis (Figure 1E) have not been explored.

**Figure 1.**
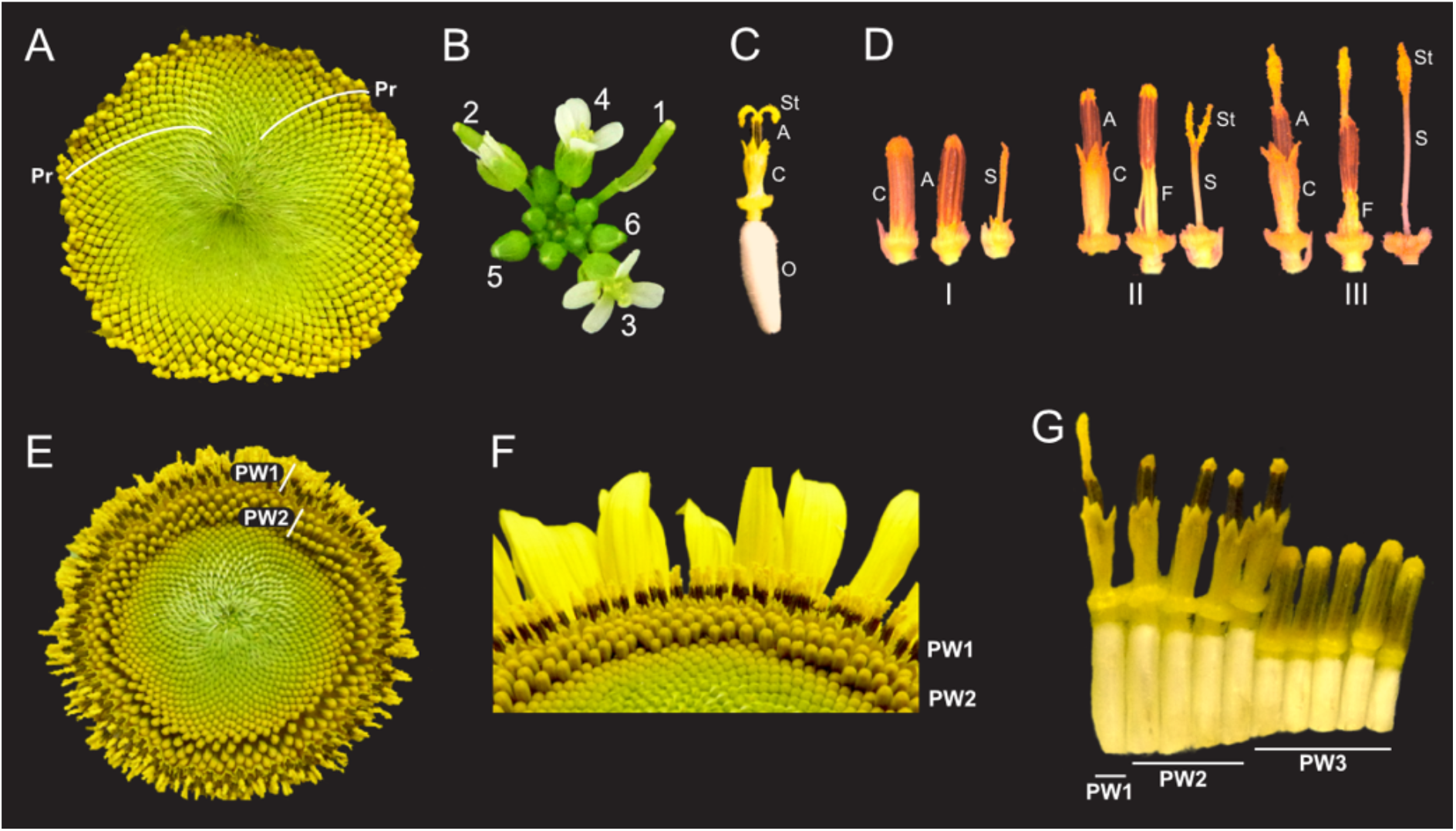
Architecture of the Sunflower Capitulum. **(a)** A sunflower capitulum before onset of anthesis, with obvious clockwise and counter-clockwise parastichies (Pr) of individual florets. **(b)** An *Arabidopsis thaliana* inflorescence also displays spiral phyllotaxy. Numbers indicate age of flowers. **(c)** One sunflower floret post-anthesis. Organs are abbreviated as ovary (O), corolla tube (C), anther tube (A), filaments (F), style (S), and stigma (St). Anther tubes and filaments collectively make the stamens. **(d)** The primary stages of floret development: (I) pre-anthesis immature, (II) early anthesis staminate, and (III) late anthesis pistillate. For each stage, the left-most floret is undissected, the corolla tube was removed from the middle floret, and the corolla and stamen were removed from the right-most floret. **(e)** A sunflower capitulum on day two of anthesis in which coordinated pseudowhorls (PW) of developing florets are apparent. **(f)** An angled view of **(e). (g)** A cross-section of **(e**); florets and ovaries along a single parastichy that belong to three different pseudowhorls. Plants grown in LD|25°C (16h:8h), photograph taken at ZT 2.

Here, we investigate the timing of floral organ development in various environmental conditions. We find that temperature-compensated daily rhythms in development of both male and female floral organs, and the formation of pseudowhorls, persist when flowers are maintained in constant environmental conditions in the absence of light. However, both daily rhythms in development and the formation of pseudowhorls are lost in constant light, with late-stage floret maturation occurring continuously along parastichies. Importantly, the effects of light on floret development are time-of-day-dependent. Together, these data indicate that late-stage floret development is regulated by the circadian clock. We propose that pseudowhorls are produced in response to an external coincidence mechanism, in which developmentally competent florets undergo reproductive transitions only when they receive environmental cues at the appropriate time of day. Our data thus suggest a role for the circadian clock in the spatial as well as temporal control of sunflower development.

## Results

### During anthesis the pattern of development changes from continuous to discrete

The phyllotactic spirals of florets, or parastichies, on sunflower capitula represent continuous age-gradients with the first-specified florets on the outside edge of the capitulum (Figure 1A) (Marc and Palmer, 1981). Each disc floret is an individual flower with an epigynous ovary that holds a corolla tube of fused petals (Figure 1C). Within the corolla tube is the fused anther tube, subtended by five filaments, collectively forming the stamen (Figure 1C). Finally, at the center of the floret is the style and stigma (Figure 1C), which we will together call the style. Although early floret development occurs continuously across the capitulum, anthesis does not. Local synchronization of floret development at this developmental stage converts the obvious spiral arrangement of florets on immature capitula into a ring-like pattern of pseudowhorls (Figure 1A vs Figure 1E).

Previous studies described daily patterns in sunflower floret anthesis, with a single floret releasing pollen the day before the stigma of this floret is receptive to pollen (Putt, 1940; Neff and Simpson, 1990). Before anthesis all reproductive organs are enclosed within the corolla tube (Figure 1D – I). As anthesis begins, the corolla tubes of multiple florets within a pseudowhorl coordinately rise above the capitulum surface (Figures 1E, 1F) driven by ovary growth rather than expansion of the corolla (Figures 1G, 2A). Next, rapid elongation of anther filaments causes anther tube protrusion above the corolla (Figure 1D – II) (Baroncelli et al., 1990; Lobello et al., 2000). In this staminate stage of anthesis, pollen is released within the anther tube and is then pushed up and out by the rapid growth of the style through the anther tube soon after dawn (Figure 1D – III). Continued style elongation pushes the stigma beyond the anther tube to initiate the pistillate stage of anthesis (Figure 1D – III). This plunger-style mechanism of pollen presentation is an ancestral feature of the Asteraceae family (Jeffrey, 2009). The stigmas then unfold and become receptive to pollen the day after pollen release (Figure 1C). The temporal and spatial separation between pollen release and stigma presentation in sunflowers promotes cross-fertilization across head (Patterson, 2009).

### Pseudowhorls are generated by synchronized, daily rhythms in floret development

To better understand the synchronization of anthesis within each pseudowhorl, we characterized the detailed temporal dynamics of late-stage floret development. Plants were maintained in light-dark cycles and constant temperature. Florets within a single pseudowhorl were dissected every 15 – 30 minutes over 24 hours and individual organ lengths measured. Ovary elongation was observed first, starting near the time of lights off and continuing for approximately 10 hours (Figure 2A). As previously described (Baroncelli et al., 1990; Lobello et al., 2000), we found that stamen elongation commenced approximately 8 hours after lights off, driven by an approximately 2.5x increase in anther filament length over 5 hours (Figure 2A). Anther tubes, like corolla tubes, did not elongate during this time frame. Stamens reached maximal length at time 28, and later shrank due to filament collapse (Figure 2A, Figure 1D – III). Styles initiated rapid elongation one hour after anther filaments, with growth continuing over 11 hours until they reached ∼2x their original length and grew past the stamens. Thus, the obvious discrete phases of anthesis visible on whole capitula (Figures 1E – G) are driven primarily by the consecutive and highly coordinated elongation of ovaries, anther filaments, and styles within each pseudowhorl.

**Figure 2.**
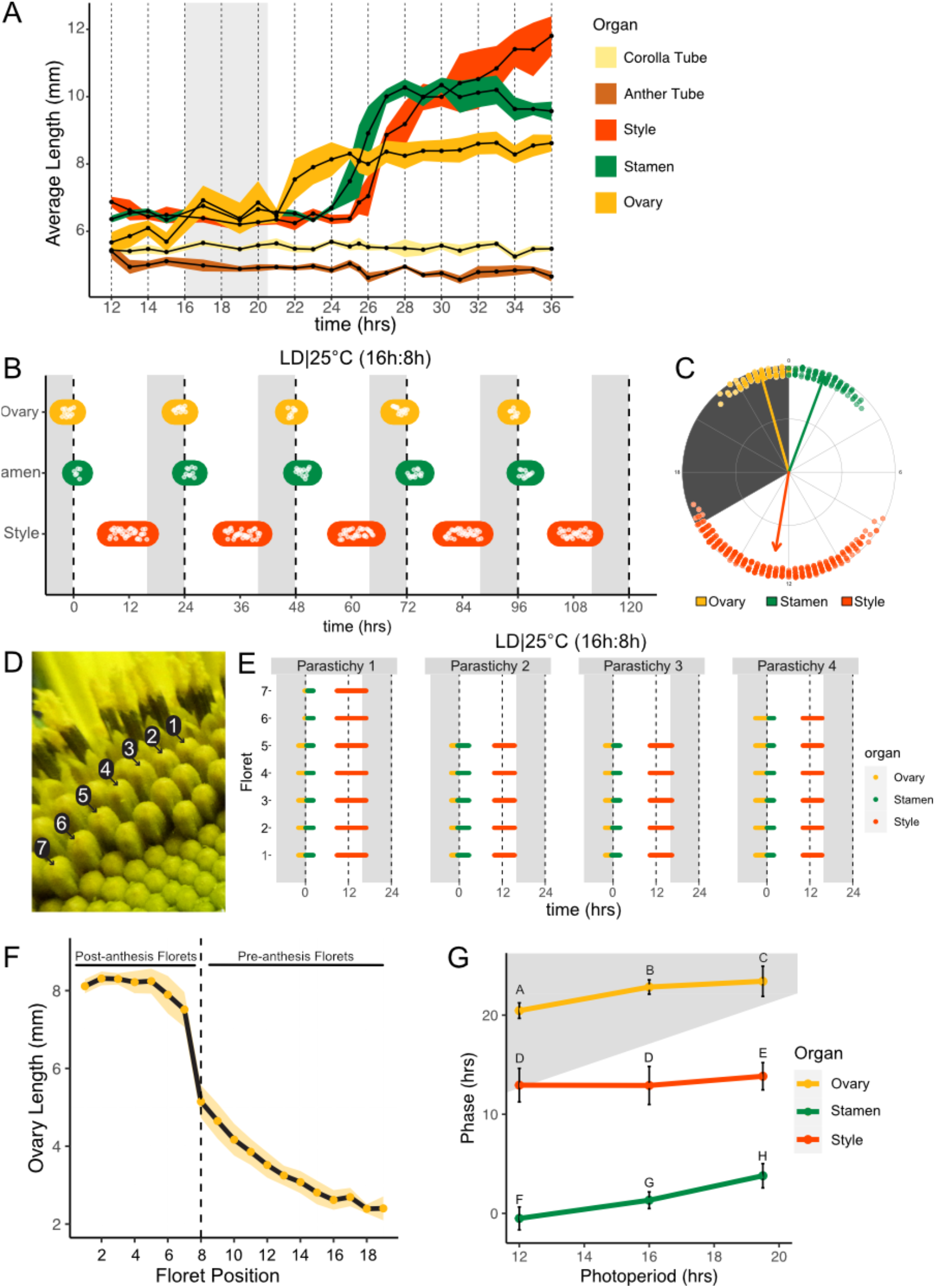
Pseudowhorls are Due to Coordinated Daily Rhythms in Floret Anthesis. **(a)** Growth kinetics of floral organs in a single pseudowhorl for plants grown in LD|25°C with dark from ZT 16-20.5. Corolla tube, anther tube, style + stigma, stamen, and ovary were measured. Black dots represent average lengths and ribbons represent standard deviation. (n = 7 – 15 florets per time point; n = 1 biological replicate) **(b)** Timing of ovary, stamen, and style growth for florets on a capitulum in LD|25°C (16h:8h). Each white point represents time when organs of >5% of florets per pseudowhorl elongated (n=3 capitula); colored bars group growth of organs within a pseudowhorl. **(c)** The timing of active organ growth for pseudowhorls as seen in **(b)**, superimposed on a 24-hour circular clock (n=3 capitula). Dots represent individual observations, arrow directions represent average phases, and arrow lengths represent precision of timing. **(d)** Numbered florets within a single parastichy. **(e)** Timing of active growth for all florets along a parastichy within pseudowhorl 2; plants grown in LD|25°C (16h:8h) (n = 4 parastichies). **(f)** Lengths of ovaries along a parastichy at ZT 2; plants grown in LD|25°C (16h:8h). Position 8 represents the first floret in the next pseudowhorl (n=4 parastichies). Yellow dots indicate means and the ribbon represents standard deviation. **(g)** The average phases of active growth for ovaries, stamens, and styles in LD|25°C with dark from either ZT 12-24 (left), ZT 16-24 (middle), or ZT 16-20.5 (right); (n = 3-4 capitula). Error bars are standard deviation. Different letters indicate statistically significant differences in mean phases by one-way ANOVA with post-hoc Tukey HSD test (*P* < 0.001). For all graphs the period of dark exposure is shown with grey background.

We next investigated the effects of different environmental conditions on the dynamics of floret development. We used time-lapse movies of sunflowers undergoing anthesis taken at 15-minute intervals and scored observable patterns of growth as proxies for the detailed organ measurements described above. We first examined growth kinetics in capitula undergoing anthesis in long-day conditions reminiscent of a summer day (16 hours light, 8 hours dark, and constant 25°C (LD|25°C (16h:8h)). A new pseudowhorl of florets underwent anthesis every 24 hours, with ovary swelling initiating in the late night, stamen elongation visible from late night to soon after dawn, and conspicuous style elongation occurring in the latter part of the day and early night (Figure 2B; Supplemental Table 1; Video 1). Times of ovary, stamen, and style growth were highly coordinated across pseudowhorls and flowers (Figure 2C). The mean midpoint of visible ovary growth occurred before dawn at Zeitgeber Time (ZT) 22.9 (with the time of lights on defined as ZT 0), followed by the mean midpoint of stamen growth at ZT 1.3, and the mean midpoint of style growth before dusk at ZT 12.6 (Figures 2B, 2C; Supplemental Table 1). Similar experiments were carried out for plants exposed to shorter (LD|25°C (12h:2h)) or longer (LD|25°C (19.5h:4.5h)) photoperiods. Similar daily rhythms of anthesis were observed in all three conditions, with the phases of organ growth relative to dawn changing only slightly with different night lengths (Figure 2G; Supplemental Figure 1). In all three photoperiod conditions, the timing of floret development within pseudowhorls was highly coordinated (Figure 2B; Supplemental Figure 1).

This high degree of developmental coordination within each pseudowhorl is quite remarkable given that the initiation of floret primordia along a parastichy occurs continuously over many days (Zhang et al., 2021). We therefore examined the timing of ovary, stamen, and style growth within a pseudowhorl along separate parastichies (Figures 2D, 2E). While the number of florets per parastichy in a pseudowhorl varied, growth of the three different organs occurred at almost the identical time for all florets (Figure 2E). Ovary lengths were also measured along individual parastichies at ZT 2, a time when ovary growth within a pseudowhorl is complete (Figure 2A). The lengths of ovaries within a pseudowhorl along a parastichy were similar to each other (Figures 1G, 2F), but there was a large step-down in length compared to ovaries of pre-anthesis florets in the same parastichy. Previous studies showed the large changes in ovary size and filament length during anthesis are due to rapid cell expansion (Lindström et al., 2007; Lindström and Hernández, 2015; Lobello et al., 2000). The remarkable synchronization of cell elongation and organ growth during anthesis, despite the age difference of florets along each parastichy, suggests some master regulator coordinates anthesis within each pseudowhorl.

### Daily rhythms in floret development persist in constant dark conditions and are temperature compensated

The precise daily timing of sunflower anthesis suggested the circadian clock might be involved in this process. To test this, we examined the development of sunflower capitula maintained in constant dark and temperature. Rhythms in ovary and stamen elongation persisted in these free-running conditions, with a new pseudowhorl of florets undergoing anthesis every ∼24 hours with phases similar to those seen in light/dark cycles (Figures 3A, 3B; Video 2). Scoring of individual florets along a parastichy confirmed coordinated floret organ growth within a pseudowhorl (Figure 3C), even after 4 days in DD|25°C (Supplemental Figure 3). We next examined ovary lengths among the florets along individual parastichies. As in photoperiodic conditions, ovary lengths between pseudowhorls were clearly distinct (Figures 3E, 3F). Thus, daily rhythms in anthesis and the formation of pseudowhorls persisted in the absence of environmental cues, suggesting regulation by the circadian clock.

**Figure 3.**
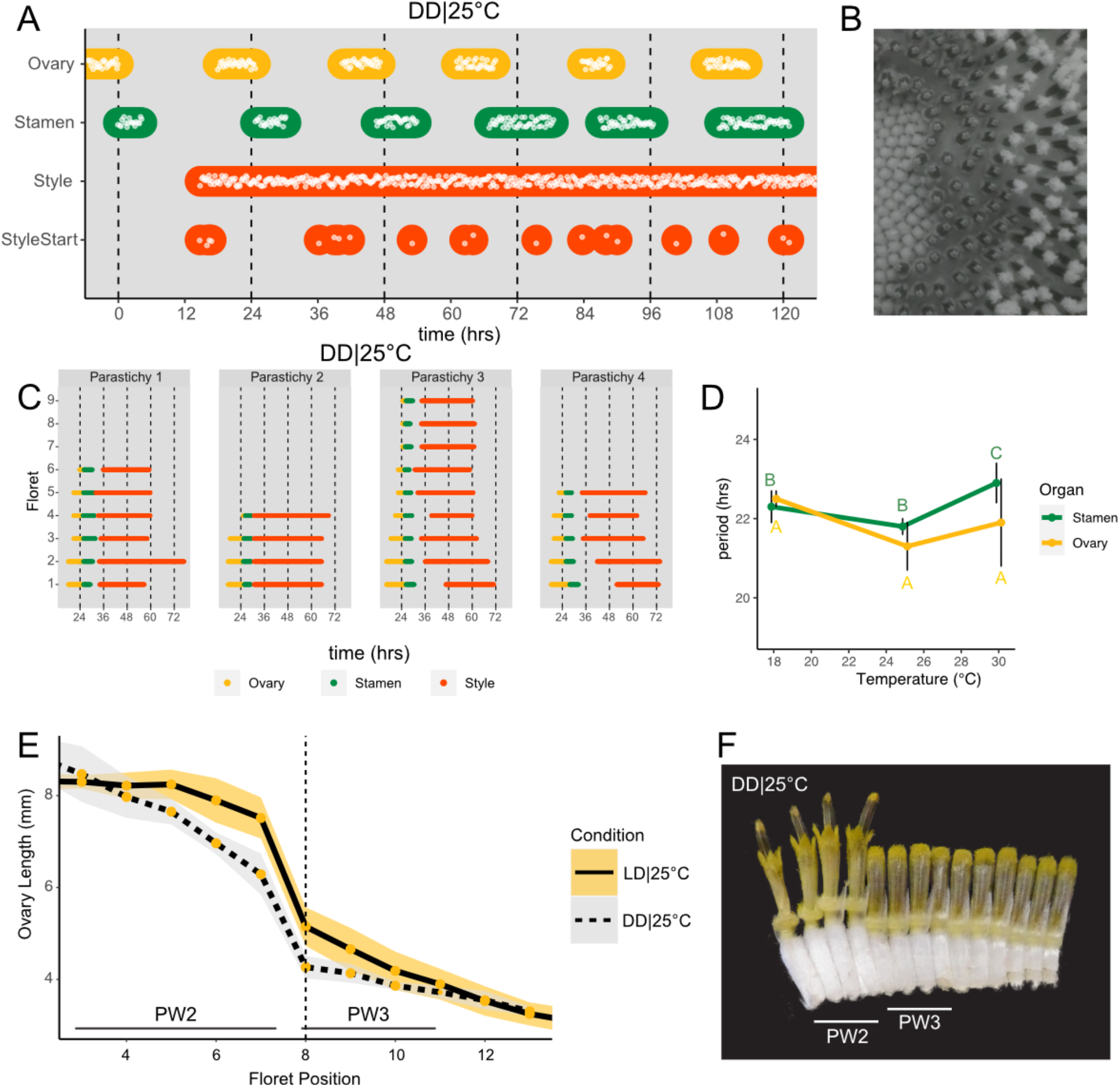
Daily Rhythms in Floret Anthesis are Regulated by the Circadian Clock. **(a)** Timing of ovary, stamen, style, and start-of-style growth for florets in DD|25°C (n=4 capitula). **(b)** Sunflower capitulum in DD|25°C at ZT 50, showing coordinated floret anthesis. **(c)** Timing of active growth for all florets per parastichy within pseudowhorl 2; DD|25°C (n = 4 parastichies). **(d)** Average periods (hrs) of rhythmic ovary and stamen growth in DD|18°C (n=3), DD|25°C (n=4), or DD|30°C (n=3 capitula). Error bars represent standard deviation. Different letters indicate statistically significant differences by one-way ANOVA with post-hoc Tukey HSD test (*P* < 0.001), with yellow letters for differences within ovaries and green letters for differences within stamens. Periods of stamen and ovary growth are not statistically different within any temperature treatment. **(e)** Lengths of ovaries along a parastichy for sunflowers grown in either LD|25°C (16h:8h) (n=4 parastichies) or DD|25°C (n=8 parastichies) at ZT 26, with position 8 the first ovary in the next pseudowhorl. Yellow dots indicate means and the ribbons standard deviation. **(f)** A cross-section of a capitulum grown in DD|25°C; florets and ovaries along a single parastichy that belong to two different pseudowhorls. Photograph taken at ZT 26. Plots as described for Figure 2.

As seen for ovaries and stamens, the start of visible style elongation within each pseudowhorl occurred at the expected times and was highly coordinated for the first two days in constant darkness (Figures 3A, 3C; Supplemental Figure 3). However, style elongation rates were very slow in DD|25°C (Supplemental Figure 4) such that style growth in one pseudowhorl continued even after florets in the subsequent pseudowhorl began anthesis. This resulted in almost continuous style elongation across the capitulum. The immediate slowing of style but not ovary or stamen elongation upon transfer to constant darkness (Supplemental Figure 4) suggests that different regulatory pathways control growth of these organs. However, the rhythmic initiation of growth of all three types of organs in constant darkness suggests a general role for the circadian clock in the control of anthesis.

**Figure 4.**
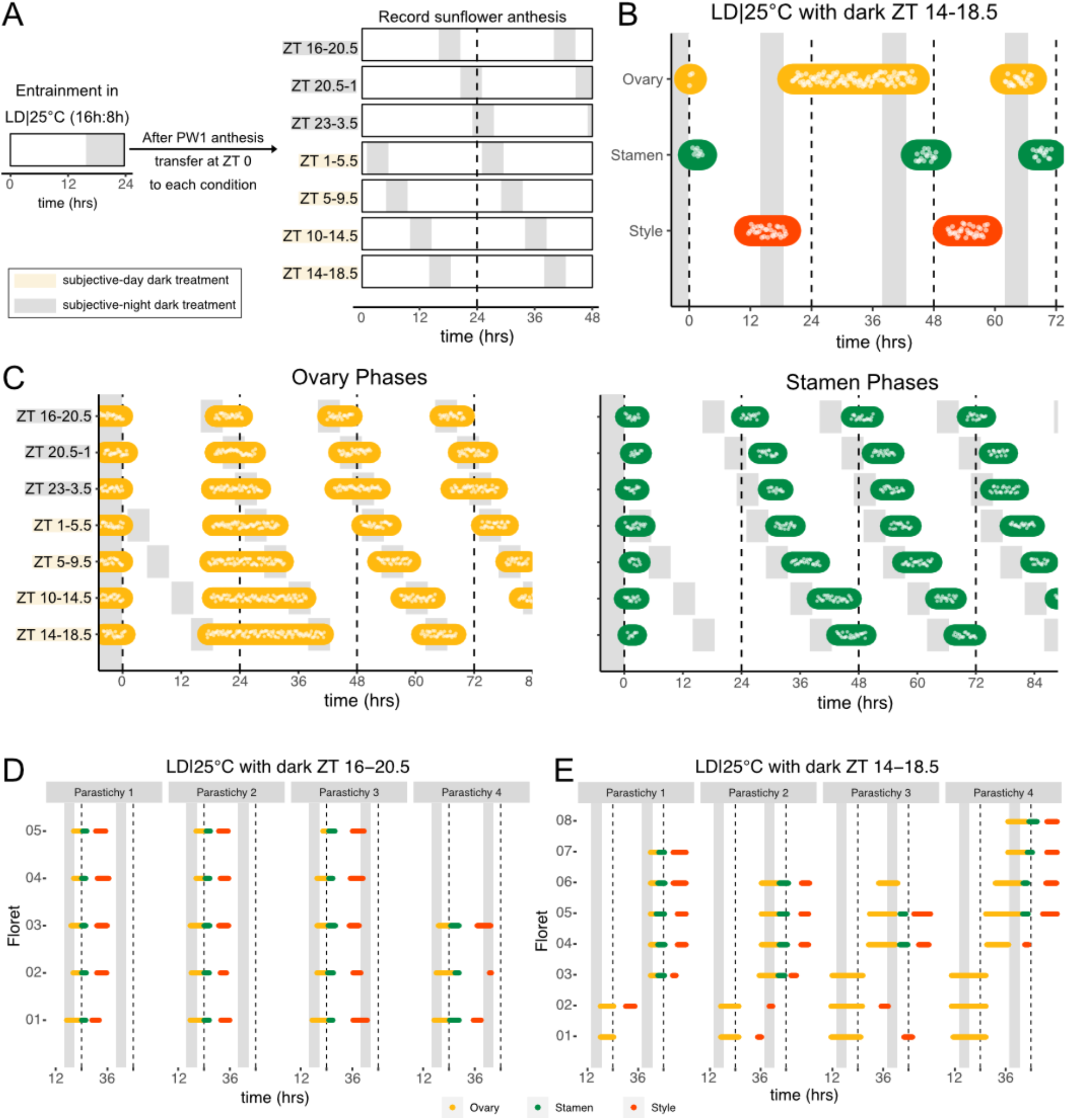
Normal Rhythms of Anthesis are Maintained with Darkness During the Subjective Night but Not During the Subjective Day. **(a)** Schematic representation of experiment. All plants were maintained at 25°C with 4.5 hr of darkness provided at the times indicated with grey boxes **(b)** Timing of ovary, stamen, and style growth with dark during ZT 14-18.5 (n=4 capitula). **(c)** Timing of ovary (left) and stamen (right) growth with darkness provided during the indicated times (n=3-4 capitula per condition). **(d, e)** Timing of active growth for all florets along a parastichy within pseudowhorl 2 of sunflowers with dark during **(d)** ZT 16-20.5 or **(e)** ZT 14-18.5 (n = 4 parastichies for each condition). Plots are as described for Figure 2.

Circadian-regulated processes are characterized by their relative insensitivity to temperature variation across a physiologically relevant range (Greenham and McClung, 2015). We therefore determined the free-running periods for ovary and stamen developmental rhythms in sunflowers maintained in constant darkness at 18°C, 25°C, or 30°C. Across the three different growth conditions, the estimated periods for cycles of ovary development were not statistically different while the estimated period for stamen development was slightly but significantly longer at 30°C, suggesting modest overcompensation (Figure 3D; Supplemental Table 1). However, within each temperature, the estimated periods of ovary and stamen growth were not significantly different (Figure 3D). The rhythmic internal processes that regulate timing of floret anthesis are therefore temperature compensated, further suggesting regulation by the circadian clock.

### The circadian clock gates floral responsiveness to dark cues

Environmental influences on clock-regulated processes are often ‘gated’ by the circadian clock, meaning that the same stimulus can evoke different responses when applied at different times of day (Wenden et al., 2011). We therefore examined whether the effects of light and darkness on sunflower anthesis were time-of-day dependent by applying a short 4.5-hour dark period at different times of the subjective day or night (Figure 4A) and assessing the effects on the spatial and temporal regulation of floret development. When the onset of darkness occurred during the subjective night, anthesis proceeded in the expected pattern with minor phase adjustments (Figures 2A, 4C; Supplemental Figure 5). However, when the onset of darkness occurred during the subjective day, different effects were observed for ovaries compared to stamens and styles. In pseudowhorl 2, the first pseudowhorl undergoing development in the new light conditions, ovary elongation began at the expected time but with a slow growth rate that continued until the next period of darkness (Figures 4B, 4C; Supplemental Figure 5). In contrast, development of stamens and styles did not initiate until ovary growth within pseudowhorl 2 was complete. As a result, dark treatment during the subjective day caused a considerable lag in anthesis (Figure 4C; Supplemental Figure 5). After two days in each new photoperiodic condition, the phases of ovary, stamen, and style elongation shifted such that maturation of all floret organs occurred with stable phase relationships to the new times of dusk and dawn (Supplemental Figures 5A-G, 5I). This rapid shift in the phase of anthesis suggests re-entrainment of the circadian clock may occur quickly in sunflower capitula.

**Figure 5.**
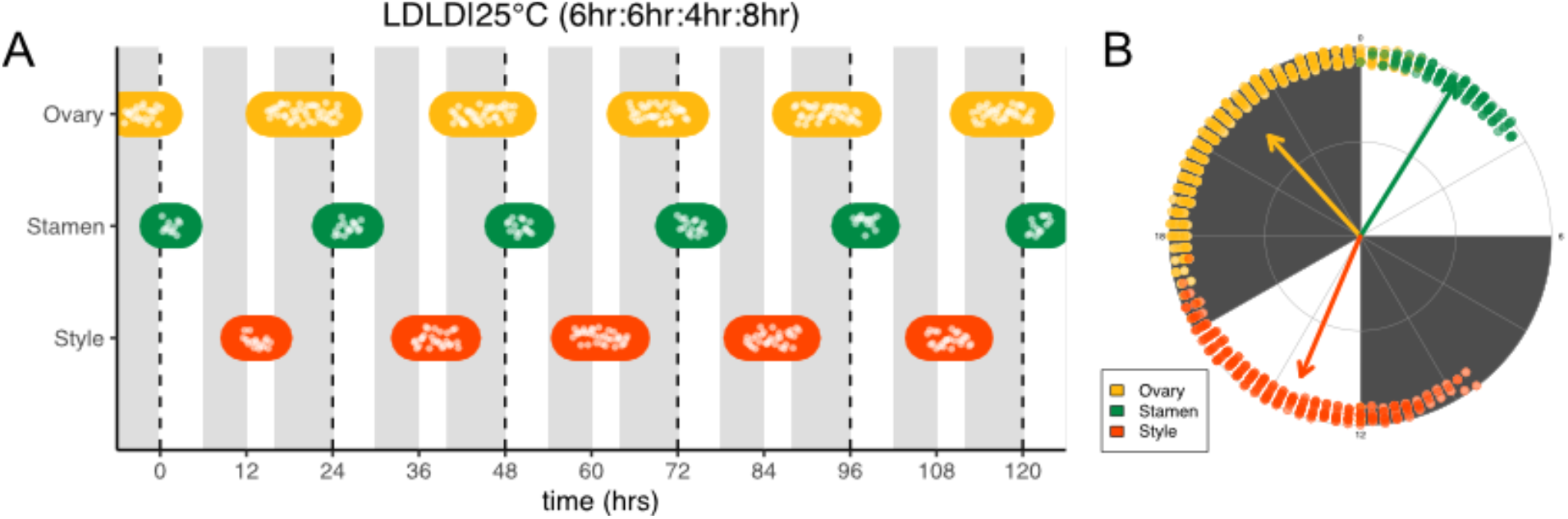
Frequency Demultiplication of Sunflower Anthesis Rhythms. **(a)** Timing of ovary, stamen, and style growth for florets on a capitulum with two cycles of light and dark each day, namely 6 hrs L, 6 hrs D, 4 hrs L, 8 hrs D (n=3 capitula). **(b)** The timing of active organ growth for pseudowhorls as seen in **(a)**, superimposed on a 24-hour circular clock (n=3 capitula). Plots as described for Figure 2.

We further characterized anthesis in the ZT 14-18.5 condition, in which darkness occurred two hours earlier than in the original entraining conditions. In this condition, there was a 42.9-hour difference between start of pseudowhorl 2 ovary growth and the start of pseudowhorl 3 ovary growth, compared to the 22.8-hour difference seen in the ZT 16-20.5 dark condition (Figures 4B, 4C; Supplemental Figure 5J, Supplemental Table 2). Surprisingly, this large delay in anthesis did not cause two pseudowhorls to develop simultaneously: the average number of new florets undergoing anthesis within the new pseudowhorl stayed the same across the different light treatments (Supplemental Figure 6). Close inspection of the timing of development of individual florets along parastichies within pseudowhorl 2 revealed desynchronization of anthesis in the ZT 14-18.5 dark condition (Figure 4E). In contrast, dark treatment from ZT 16 – 20.5 promoted the highly coordinated pattern of anthesis seen in LD|25°C (16h:8h) conditions (Figures 2E, 4D). The large difference in maturation patterns depending upon whether lights were turned off at ZT 14 or at ZT 16 demonstrates a time-of-day dependent response to dark cues, a further indication of circadian clock regulation of this process.

**Figure 6.**
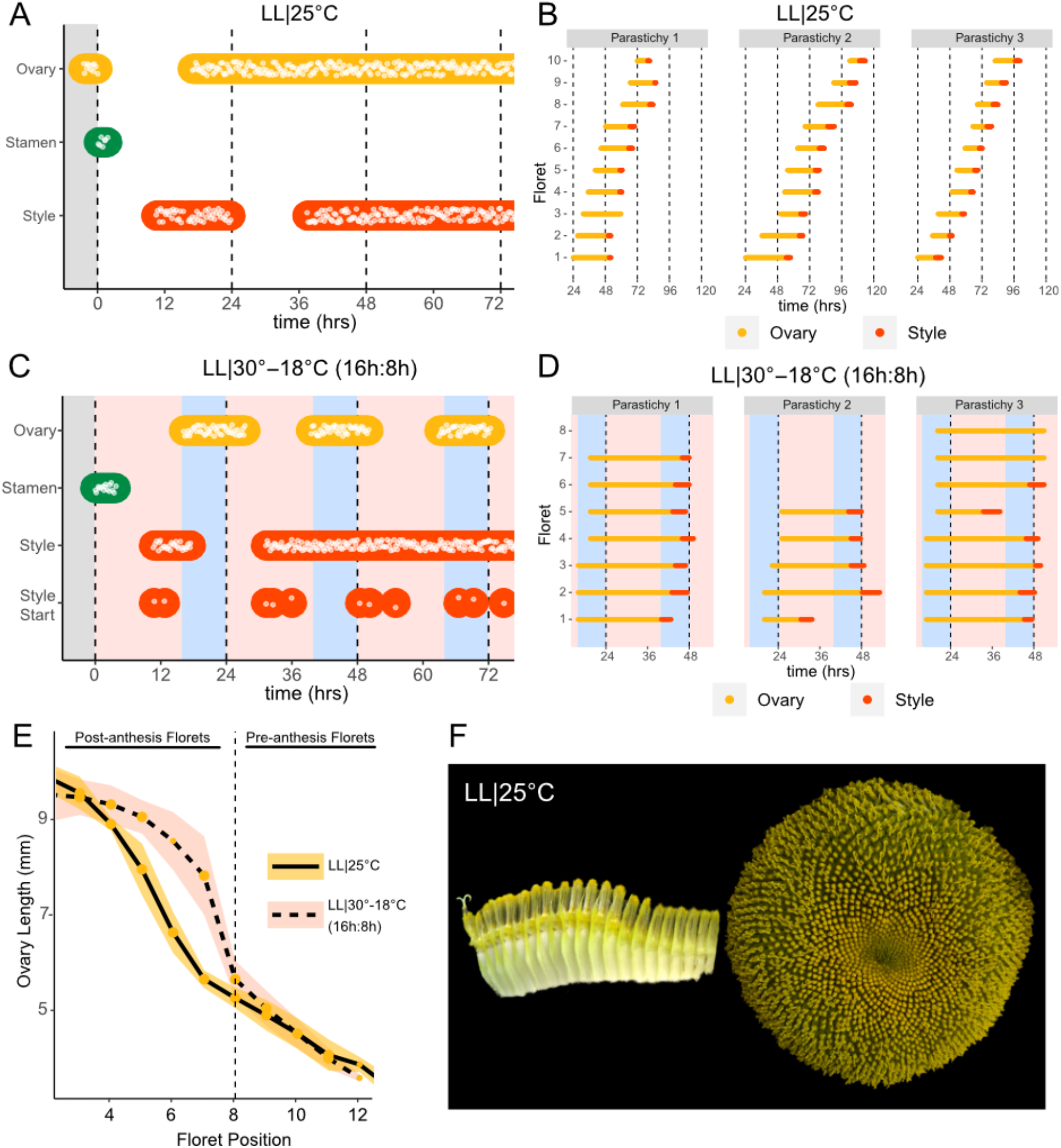
Constant Light Disrupts Coordination of Anthesis and Pseudowhorl Formation. **(a, c)** Timing of ovary, stamen, and style growth for florets on a capitulum in **(a)** LL|25°C (n=3 capitula), and **(c)** LL|30°-18°C (16h:8h) (n=3 capitula). **(b, d)** Timing of active growth 24 hours after transfer to **(b)** LL|25°C, 10 consecutive florets measured along each parastichy (n = 3 parastichies), and **(d)** LL|30°-18°C (16h:8h) for all florets per parastichy in pseudowhorl 2 (n = 3 parastichies). **(e)** Lengths of ovaries along a parastichy for sunflowers grown in either LL|25°C (n=4 parastichies) or LL|30°-18°C (16h:8h) (n=5 parastichies) at ZT 26, with position 8 the first ovary in the next pseudowhorl. **(f)** Florets along a parastichy at ZT 26 and the comparable sunflower capitulum at ZT 120; plants grown in LL|25°C. Periods of constant light at 30°C (pink background), 25°C (white background), and 18°C (blue background) are indicated; plots otherwise as described for Figure 2.

Many circadian-regulated processes exhibit frequency demultiplication; that is, they maintain 24-hour rhythms even when subjected to light/dark cycles with a cycle length close to half that of the endogenous period (Roenneberg et al., 2005). We therefore examined whether multiple periods of darkness during a 24-hour period would affect the timing of floret anthesis within pseudowhorls. We first exposed sunflowers to four equal periods of darkness and light every 24 hours, LDLD|25°C (6h:6h:6h:6h). One pseudowhorl of florets underwent anthesis every 24 hours, but florets within each pseudowhorl were poorly coordinated (Supplemental Figures 2K, 7B).

We next changed the proportion of darkness in a 24-hour period to a 6-hour period of darkness during the subjective day followed by an 8-hour period of darkness during the subjective night (LDLD|25°C (6h:6h:4h:8h). This combination restored coordination of 24-hour rhythms of floret development within each pseudowhorl (Figure 5A). In addition, phases of organ development were very similar to those observed in LD|25°C (16h:8h) entrainment conditions (Figure 5; Supplemental Figures 7A, 8B; Supplemental Table 1). Thus, the 6-hour dark period during the subjective day had no effect on the timing of anthesis, further suggesting circadian clock gating of responses to darkness.

**Figure 7.**
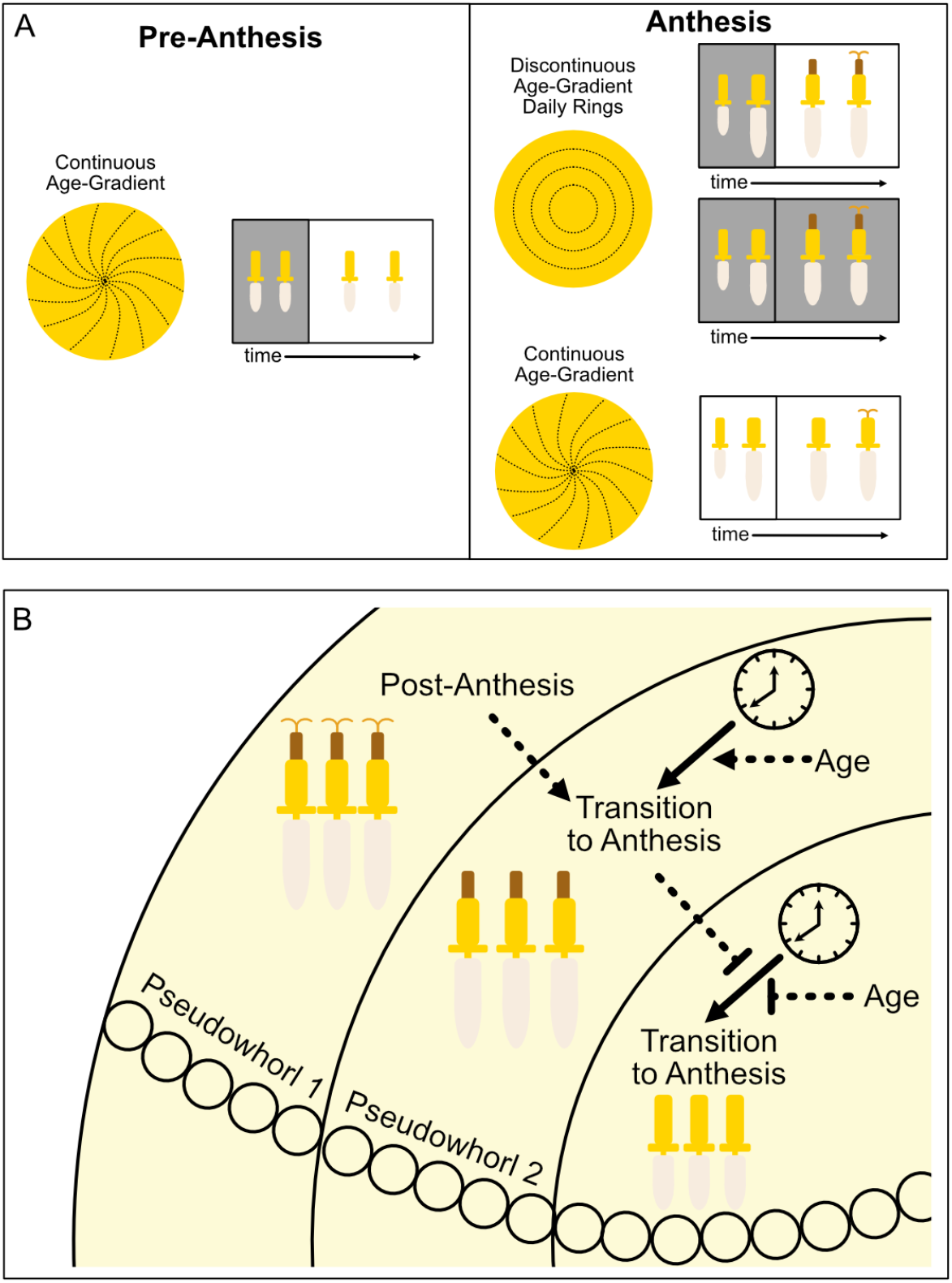
Model of sunflower anthesis patterns. **(a)** Developmental patterns observed when sunflowers undergo anthesis in different environmental conditions. Pre-anthesis, the apparent pattern of florets on a capitulum is a continuous age-gradient with spiral phyllotaxy (left panel). During anthesis in both normal light/dark cycles and in constant darkness, the capitulum transitions to a discontinuous age-gradient of florets in which rings of florets, or pseudowhorls, undergo synchronized daily rhythms of development (top right panel). In constant light conditions, however, the capitulum maintains a continuous age-gradient of floret development. Ovaries and styles grow, but stamens do not (bottom right panel). **(b)** Speculative model for transition of floret development from age-dependent continuous gradients to organization into discrete pseudowhorls that undergo coordinated anthesis. After anthesis, florets comprising pseudowhorl 1 send an unknown signal (dashed line) to younger florets, creating a permissive state for the transition to anthesis. In order to respond to this signal, younger florets must both 1) be developmentally competent, and 2) be exposed to a sufficient period of darkness during the subjective night (as determined by the circadian clock). Florets younger than pseudowhorl 2 do not elongate floral organs until the post-anthesis state of pseudowhorl 2, floret developmental status, environmental cues, and the circadian clock allow transition to anthesis and the generation of the next pseudowhorl.

### Pseudowhorl formation depends upon coordinated daily rhythms of anthesis

Previous studies show that sunflower anther filament elongation is completely inhibited and style elongation is reduced in constant light conditions (Baroncelli et al., 1990; Lobello et al., 2000). To investigate whether constant light also affected other aspects of flower development, we entrained sunflowers in LD|25°C (16h:8h) and then transitioned them to constant light and temperature (LL|25°C) on the first day of anthesis. As expected, no stamen growth was observed (Figures 6A, 6F; Supplemental Figure 9A; Video 3). Ovaries and styles did elongate, but initiation occurred continuously across the capitulum. Instead of the coordinated floret anthesis seen along a parastichy of florets in LD|25°C (16h:8h) (Figure 2E), in LL|25°C florets showed a continuous, age-dependent pattern of anthesis (Figures 6A, 6B; Supplemental Figure 9B; Video 3). Consistent with continuous rather than discrete developmental transitions across the capitulum in constant light, ovary length along a parastichy showed a size gradient dependent on the age of the floret (Figures 6E, 6F). Thus, in constant light conditions the timing of anthesis mirrors the continuous pattern of floret specification seen very early in capitulum development. This loss of circadian coordination of anthesis causes a loss of coordination of florets into pseudowhorls (Figure 6F).

We next assessed the effects of temperature cycles on anthesis in plants maintained in constant light or dark. In constant darkness with temperature cycles of 16 hours at 30°C and 8 hours of 18°C (DD|30°-18°C (16h:8h)), rhythms in the onset of anthesis were similar to those seen without temperature cycles albeit with enhanced coordination of initiation of style elongation in each pseudowhorl (Supplemental Figures 2I, 10A; Supplemental Table 1). We next investigated whether temperature cycles could rescue daily rhythms of floret anthesis in constant light (LL|30°-18°C (16h:8h)). Although anther filament growth was not restored, the addition of thermocycles to constant light coordinated the onset of ovary and style growth so that pseudowhorls formed (Figure 6C; Supplemental Figures 2H, 9C; Supplemental Table 1). Scoring of individual florets along a parastichy also showed that anthesis was coordinated into discrete pseudowhorls in constant light with thermocycles (Figure 6D; Supplemental Figure 9D). As expected, this generated a discontinuous change in ovary length along parastichies from post-anthesis florets in a pseudowhorl to pre-anthesis florets (Figure 6E). Thus, temperature cycles restore some daily anthesis rhythms and thereby rescue pseudowhorl formation.

## Discussion

Many aspects of floral physiology, including flower opening and scent production, vary across a day (Bloch et al., 2017). In a few cases, such as petal opening and closing in *Kalanchoë blössfedliana* and flower movements in *Nicotiana attenuata*, the circadian clock has been shown to regulate these diel rhythms (Bünsow, 1953; Yon et al., 2016). Here, we show that daily rhythms in the development of floral organs in sunflower persist in constant environmental conditions, are temperature compensated, and exhibit a time-of-day specific response to environmental cues (Figures 3A, 3D, 4, 5). We therefore conclude that the circadian clock controls floral development in sunflower to ensure release of pollen soon after dawn and the emergence of styles and stigmas late in the day. Although style elongation is largely complete near the end of the first day of floret anthesis (Baroncelli et al., 1990) (Figure 2A), stigmas do not become receptive to pollen until the following morning (Putt, 1940; Neff and Simpson, 1990). This temporal separation between release of pollen and stigma receptivity is thought to reduce self-pollination and is prevalent throughout the Asteraceae family (Jeffrey, 2009).

We found that elongation of ovaries, anther filaments, and styles are all under circadian regulation, but respond differently to environmental cues. For example, transfer to constant light caused a complete inhibition of anther filament elongation, significantly slowed ovary expansion, but had no significant effect on style elongation (Supplementary Figure 4). In contrast, transfer to constant darkness slowed style elongation considerably but had only modest effects on growth of ovaries and anther filaments (Figure 3; Supplementary Figure 4). The strong inhibition of style elongation in constant darkness may help explain why rhythms in style emergence were less robust than those of ovary and anther growth in this condition (Figure 3). Responses of the different organs to temperature were also distinct: temperature cycles slowed ovary expansion and had no effect on anther filament growth in either constant darkness or light; however, temperature cycles slowed style elongation rates in constant darkness but not constant light (Supplementary Figure 4). These results are consistent with our previous field studies that showed modest changes in ambient temperature affected style but not anther filament elongation (Creux et al., 2021). Overall, the different sensitivities of floral organs to light and temperature suggest they may be regulated by distinct growth regulatory pathways. This is consistent with studies in other plants, which revealed that different hormones regulate the growth of male and female reproductive organs (Chandler, 2011; Marciniak and Przedniczek, 2019).

While the molecular pathways controlling elongation of anther filaments and styles in sunflower are likely different, we demonstrate that their common regulation by the circadian clock results in the fast and near-synchronous release of pollen a few hours after dawn. Intriguingly, many bee- and butterfly-pollinated Asteraceae species release pollen in the morning (Budumajji and Raju, 2018; Hipólito et al., 2013; Neff and Simpson, 1990; Valentin-Silva et al., 2016) while at least one bat-pollinated member of the family releases pollen in the early night (Amorim et al., 2021). Since the precise timing of pollen release relative to dawn affects male reproductive success in sunflower (Creux et al., 2021), it is tempting to speculate that clock regulation of late-stage floret development may be widespread in the Asteraceae. Identification of the mechanisms by which the circadian system coordinates the growth of male and female floral organs to promote timely pollen release will be of great interest.

The ecological and evolutionary success of the Asteraceae family, estimated to contain ∼10% of angiosperm species, is attributed in part to their compressed inflorescences that act together as single false flowers to attract pollinators (Mandel et al., 2019). In sunflowers, the individual disk floret primordia are first initiated on the outer perimeter of the very large, flat head meristem and subsequent primordia are initiated following a centripetal pattern to generate the characteristic spiral pattern of immature florets seen across the disk (Figures 1A, 7A) (Marc and Palmer, 1981; Palmer and Steer, 1985). Recent experimental and modeling studies in gerbera daisies attributed formation of this spiral pattern to short-range interactions between spatially close initia (Zhang et al., 2021). In contrast, our work suggests that the ringlike pattern of pseudowhorls that emerges during anthesis (Figures 1E-1G, 7A) is generated by the action of the circadian clock on florets across the capitulum disk, with loss of synchronized daily rhythms in anthesis in constant light and temperature leading to loss of pseudowhorl formation (Figures 6A, 6B, 6F).

How do florets of different developmental ages temporally coordinate anthesis to generate pseudowhorls? We propose that a type of external coincidence model is at work, analogous to the mechanisms underlying photoperiodic control of the transition from vegetative to reproductive growth in many plants (Pittendrigh and Minis, 1964; Yanovsky and Kay, 2003). In our model, developmentally competent immature florets respond to a period of darkness that falls during the subjective night and is at least 4.5 hours long by undergoing the growth events that lead to anthesis (ovary expansion followed by the rapid onset of anther filament and then style elongation) (Fig. 7B). This is conceptually similar to photoperiodic control of flowering in short day plants such as rice, in which light at night inactivates a clock-regulated transcription factor that promotes the transition to flowering (Izawa et al., 2002). However, we speculate that local signals also contribute to the coordination of floret anthesis across the capitulum disk. Imposition of dark treatments during the subjective day delayed anthesis by close to 24 hours (Figure 4; Supplemental Figure 5J), until normal rhythmic patterns of anthesis resumed after entrainment to the new light conditions. However, pseudowhorls that developed subsequent to these delays contain the same number of florets as those found on plants that did not experience delays (Supplemental Figure 6). This suggests that signaling may occur between florets in adjacent pseudowhorls to control the onset of anthesis. This might be a positive cue from post-anthesis florets to promote development of the next pseudowhorl, or a negative cue from pre-anthesis florets to inhibit development of nearby florets (Figure 7B).

Plants are generally considered diurnal organisms and often have more robust circadian rhythms in constant light than in constant darkness (Michael et al., 2008). We were therefore surprised to find robust rhythms of anthesis in conditions of constant temperature and darkness but not constant temperature and light (Figures 3, 6). This is similar to conidiation rhythms in *Neurospora crassa*, which are observed in constant darkness but not in constant light (Sargent et al., 1966). An interesting question for further study is whether in constant light sunflower floret rhythms are present but masked, or whether the floral circadian clock is arrhythmic in this condition. The very rapid resetting of the different phases of organ growth to new light conditions (Supplemental Figure 5) suggests the latter may be true.

Another interesting question is why florets of different developmental ages undergo coordinated anthesis to generate pseudowhorls. It has been reported that the foraging activity of both honeybees and native bees on sunflower heads is positively correlated with the number of florets undergoing anthesis at any given time, and that seed set is strongly correlated with the number of bees foraging on heads (DeGrandi-Hoffman and Watkins, 2000; Neff and Simpson, 1990). In the case of the highly social honeybees, hive members communicate via a dance language to quickly reallocate the numbers of foragers sent to different food sources based on resource availability (Visscher and Seeley, 1982). We therefore speculate that the coordinated release of floral rewards by the hundreds of florets undergoing anthesis within a pseudowhorl each morning may be a mechanism to enhance attractiveness to pollinators and thus promote reproductive success.

## Materials and Methods

### Sunflower Floret Development Imaging and Scoring

Sunflower seeds of HA412 HO (USDA ID: PI 603993) genotype were planted into small pots of soil (Sunshine Mix #1, Sun Gro Horticulture) and germinated with a plastic lid in a PGV36 growth chamber (Conviron, Winnipeg, MB, Canada) at 25°C with 16 hours light (provided by metal halide and incandescent lamps, 300 μmol m^-2^ s^-1^) and 8 hours darkness per day. Plants were watered with nutrient water containing a N-P-K macronutrient ratio of 2:1:2. Two weeks after sowing, seedlings were transplanted to 2-gallon pots with 1 scoop of Osmocote fertilizer (SMG Brands). Approximately 60 days after sowing, sunflower capitula entering anthesis in their first pseudowhorl were transferred to an PGR15 growth chamber (Conviron, Winnipeg, MB, Canada; 200 μmol m^-2^ s^-1^ provided by metal halide and incandescent lamps) to image floret development under the indicated environmental conditions. Transfer to experimental conditions occurred at ZT 0. Sunflower stalks and capitula were taped to bamboo stakes (to avoid capitulum moving out of the camera frame). Raspberry Pi NoIR V2 cameras were mounted on Raspberry Pi 3 model B computers (Raspberry Pi, Cambridge, UK); cameras were fitted with LEE 87 75 x75mm infrared (IR) filters (Lee Filters, Andover, England). Computers were programmed to take a photo every 15 minutes. Infrared LEDs (Mouser Electronics, El Cajon, CA, USA) were programmed to flash during image capture so that the capitula were visible in the dark without disrupting plant growth. Sunflowers were imaged immediately upon transfer to experimental conditions, and through anthesis of all florets in the capitulum. The 15-minute interval images were analyzed sequentially in a stack on ImageJ (Schneider et al., 2012). For each image, the ovaries, stamens, and styles were scored for a change in size from the previous image. Ovary growth was seen as corolla tube swelling above the immature capitulum surface (Figures 1E – G). Late-stage anther filament elongation was observed starting from when the corolla tube cracked open to reveal the stamen tube until its full extension above the corolla surface (Figures 1D – II, 1G). Late-stage style elongation was observed starting with the visible extrusion of pollen out of the top of the anther tube and ending with the style fully extended (Figures 1D – III, 1G). Organs were classified as growing when >5% of the florets in a pseudowhorl showed a change in length in one time-lapse image relative to the previous one. Growth was measured qualitatively as active or inactive.

### Polar Plots and Statistical Analyses

All sunflower scoring data was plotted in R using the tidyverse package (Wickham et al., 2019). To create polar plots based on each pseudowhorl, scoring data was separated into pseudowhorls. At times, more than 24 hours of data is presented on one polar plot. In the environmental conditions that eliminated pseudowhorl coordination, polar plots were created based on 24 hours of data. The average time of anthesis for each organ, and standard deviation was calculated using the circular package in R (Agostinelli, C. and Lund, 2022). This information was calculated for each pseudowhorl, all pseudowhorls together superimposed on one 24-hour cycle, and for pseudowhorls 3 and up (after resetting is complete). The BioDare2 website (https://biodare2.ed.ac.uk/) was used to calculate rhythmic parameters for all conditions and organs (Zielinski et al., 2014). The FFT-NLLS fit method was used to calculate period and phase values based on average peak values. The time frame analyzed was different in each condition and is reported in Supplemental Table 1. R scripts used to generate each figure are available upon request.

### Scoring Consensus Growth for Florets in a Parastichy

The 15-minute interval images were analyzed sequentially in a stack on ImageJ. A clockwise parastichy was selected, and florets numbered from the outer edge of the pseudowhorl inward. At images taken at 15-minute intervals, the ovaries, stamens, and styles for each floret were scored for a change in size from the previous image. Four parastichies per condition were analyzed, from two different sunflower heads. In LL|25°C conditions, which had no pseudowhorls, 10 florets in a parastichy were scored for 48 hours. For all other conditions tested, a parastichy from the 2^nd^ pseudowhorl was scored until anthesis of all florets per parastichy within the pseudowhorl was complete. For the DD|25°C condition, the 4^th^ pseudowhorl was also scored.

### Counting Florets per Pseudowhorl

For the 4.5-hour dark conditions imposed at various times of day, the number of florets that began anthesis per pseudowhorl was counted every 24 hours. For each condition, florets from 12-19 parastichies from two different sunflower heads were counted.

### Floret Organ Timecourse and Ovary Growth Measurements

To measure the organs in florets over a timecourse (Figure 2A; LD|25°C (19.5h:4.5h)) entire florets from a single pseudowhorl were dissected from the capitulum and photographed. Organs were then measured with Image J for each floret. For ovary growth measurements, at ZT 2 on after the second day in a given growth condition, entire florets, including ovaries, were dissected from the capitulum along 4-8 continuous parastichies and photographed. Ovary length was then measured with Image J for each floret, consecutively. Position 8 always designated the boundary between pseudowhorls.

## Acknowledgements

This work was supported by grants from the National Science Foundation (IOS 1238040) and the U.S. Department of Agriculture-National Institute of Food and Agriculture (CA-D-PLB-2259-H) to SLH. We thank Chris Brooks, Hongtao Zhang, and Cassandra Baker for help tending to and running experiments on sunflowers; Julin Maloof and John Davis for helpful discussions and statistical and scripting advice; and Mike Covington for his R script to generate polar plots.

## Video Legends

**Video 1**

Anthesis on a capitulum maintained in light/dark cycles (16 hours light, 8 hours dark) at a constant 25°C.

**Video 2**

Anthesis on a capitulum maintained in constant darkness and temperature (25°C).

**Video 3**

Anthesis on a capitulum maintained in constant light and temperature (25°C).

---

**Supplemental Figure 1.**
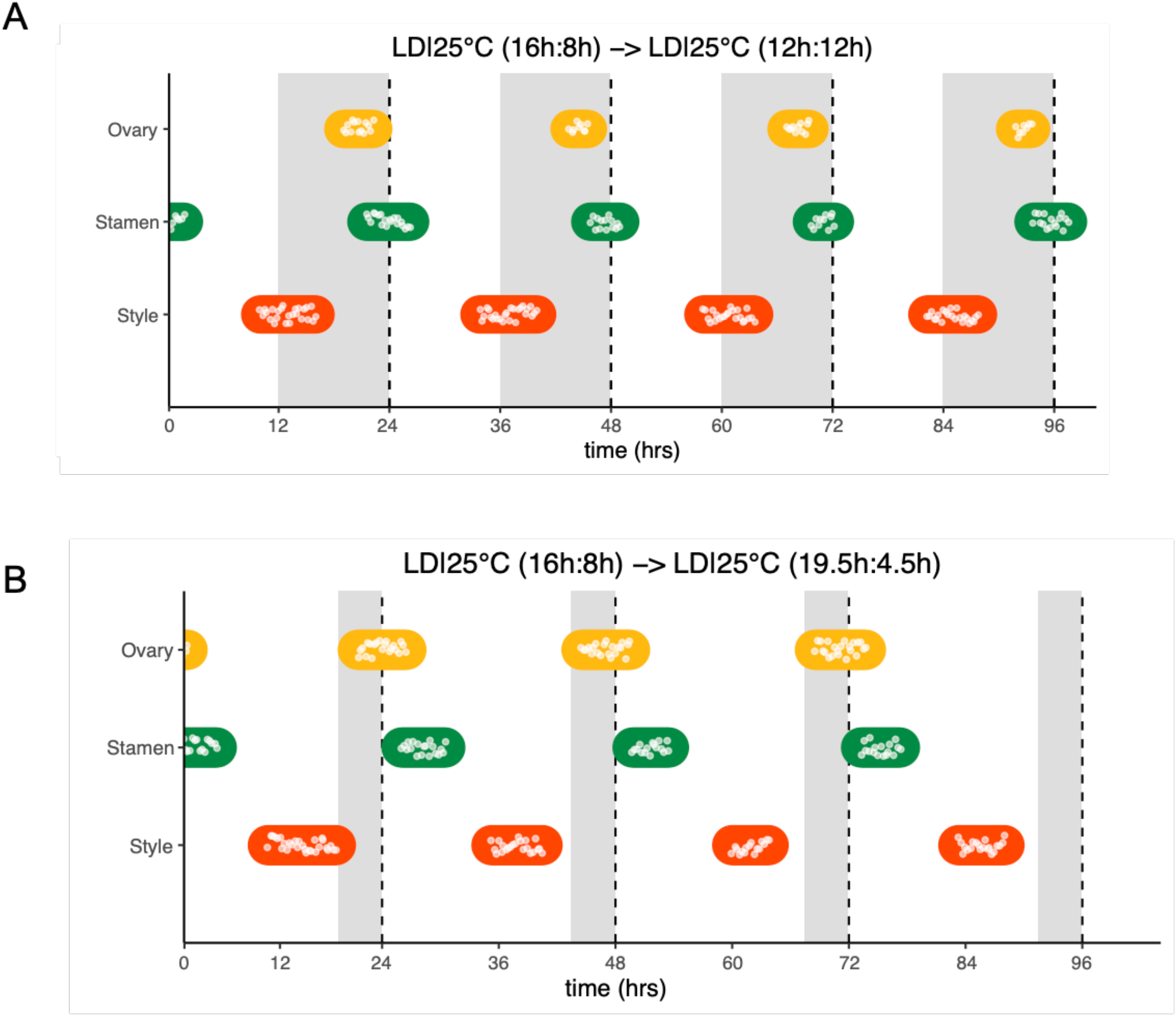
Sunflower Anthesis is Rhythmic in Different Photoperiodic Conditions. **(a, b)** Timing of ovary, stamen, and style growth for florets on a capitulum in **(a)** LD|25°C (12h:12h) and **(b)** LD|25°C (19.5h:4.5h), representing growth for >5% of florets in a pseudowhorl (n=4 capitula for each condition). Plants were grown in LD|25°C (16h:8h) and then transferred to the indicated photoperiods at time 0. White dots are individual observations of growth, and colored bars group growth for organs within a pseudowhorl. Periods of dark exposure are shown with grey boxes.

**Supplemental Figure 2.**
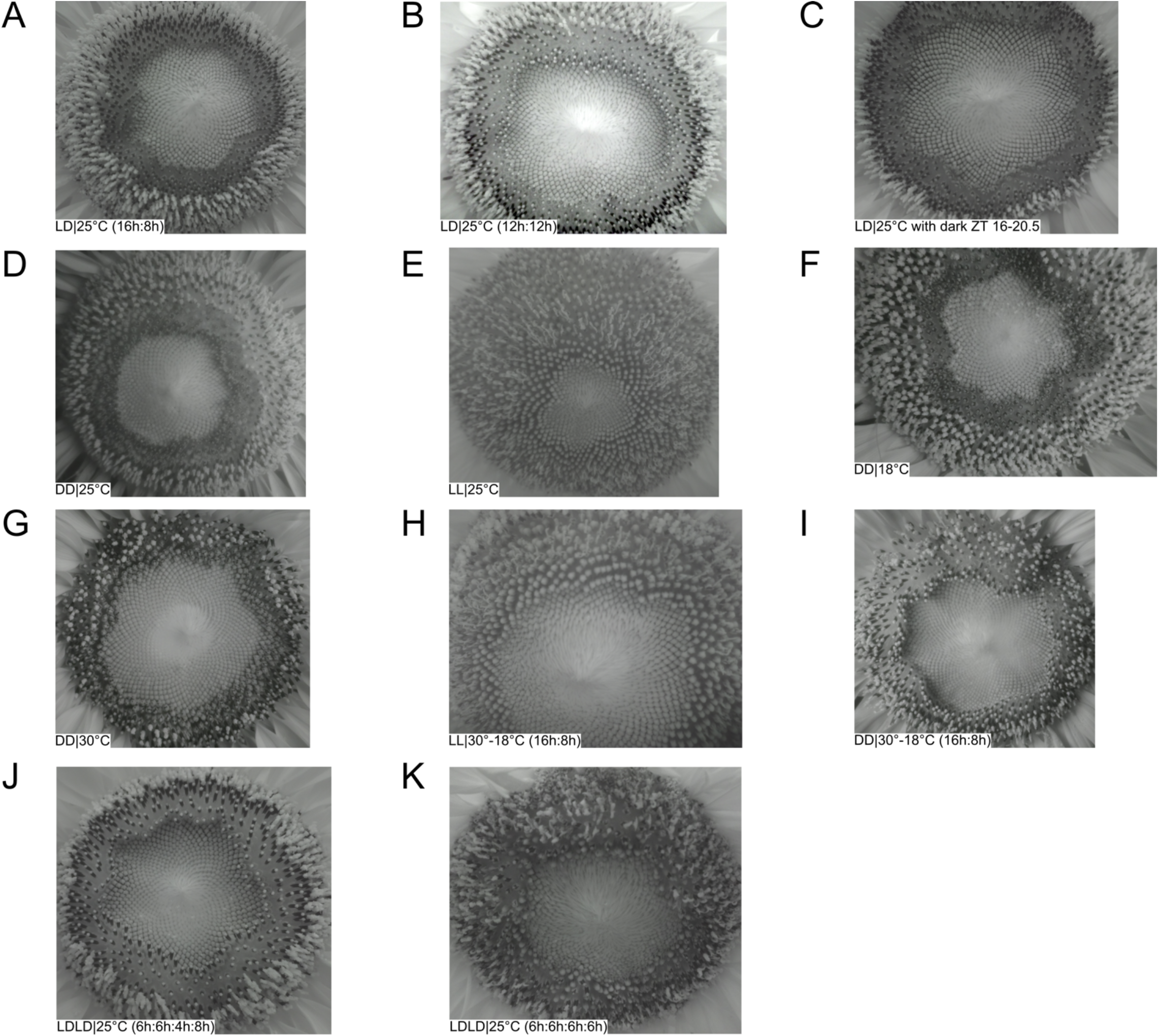
Sunflower Capitulum Development in Various Environmental Conditions. Images of sunflower capitula grown in **(a)** LD|25°C (16h:8h), **(b)** LD|25°C (12h:12h), **(c)** LD|25°C with dark ZT 16-20.5, **(d)** DD|25°C, **(e)** LL|25°C, **(f)** DD|18°C, **(g)** DD|30°C, **(h)** LL|30°-18°C (16h:8h), **(i)** DD|30°-18°C, **(j)** LDLD|25°C (6h:6h:4h:8h), and **(k)** LDLD|25°C (6h:6h:6h:6h). All images were taken 2 hours after subjective dawn relative to their entrainment conditions of LD|25°C (16h:8h), but at various numbers of days after transfer to the new environmental conditions.

**Supplemental Figure 3.**
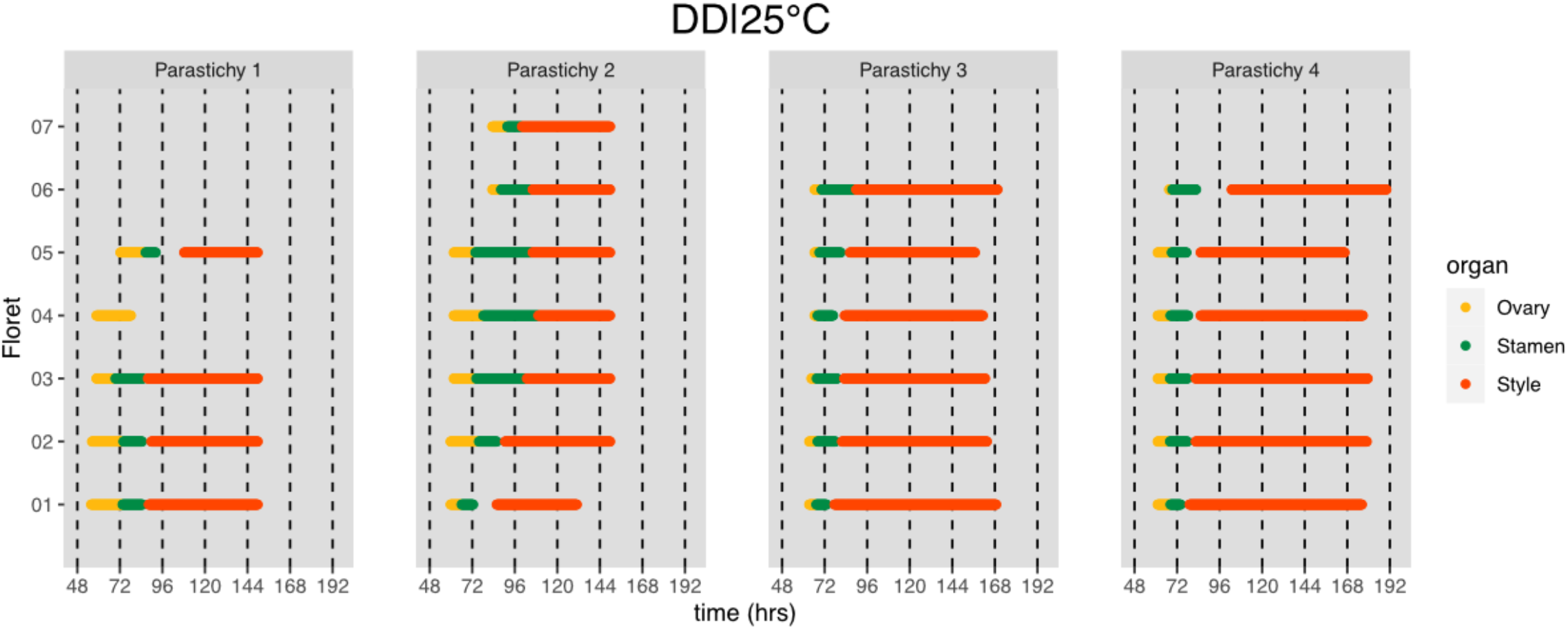
Pseudowhorl Coordination Persists for Multiple Days in Constant Dark Free-running Conditions. Timing of active growth for all florets along parastichies within pseudowhorl 4 of sunflowers after transfer to DD|25°C (n=4 parastichies).

**Supplemental Figure 4.**
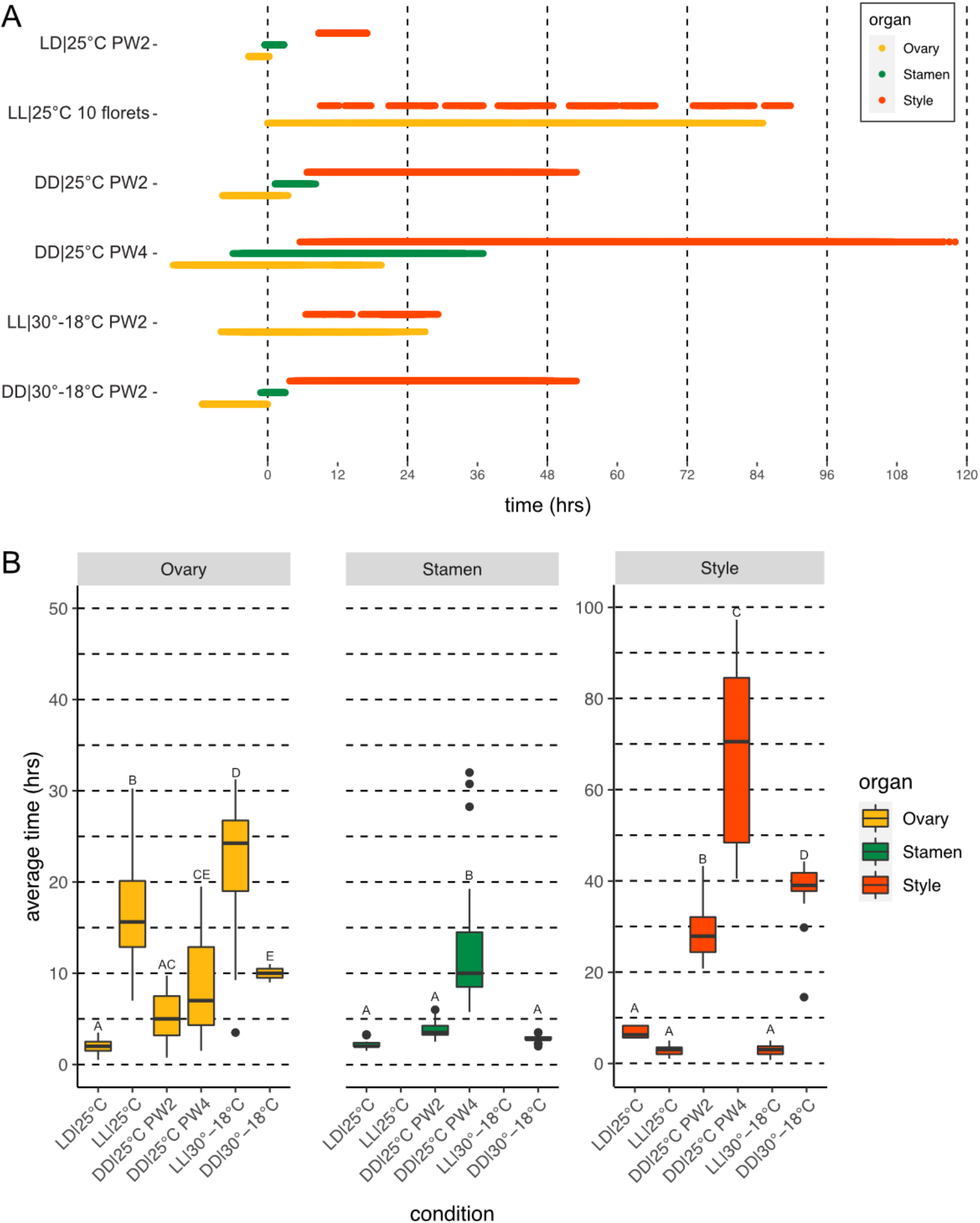
Rates of Floral Organ Maturation are Dependent on Environmental Conditions. **(a)** Total time for florets within the indicated pseudowhorls to complete anthesis, compiled from the individual floret timing for: LD|25°C (16h:8h) (Fig. 2E); LL|25°C (Fig. 6B); DD|25°C PW2 (Fig. 3C); DD|25°C PW4 (Supplemental Fig. 3); LL|30°-18°C (16h:8h) (Fig. 6D); and DD|30°-18°C (16h:8h) (Supplemental Fig. 10B). **(b)** Average time required for completion of maturation of individual florets, calculated from above data. Florets were selected from the same pseudowhorl (except in LL|25°C). Error bars indicate standard deviation. Different letters indicate statistically significant differences by one-way ANOVA with post-hoc Tukey HSD test (*P* < 0.001).

**Supplemental Figure 5.**
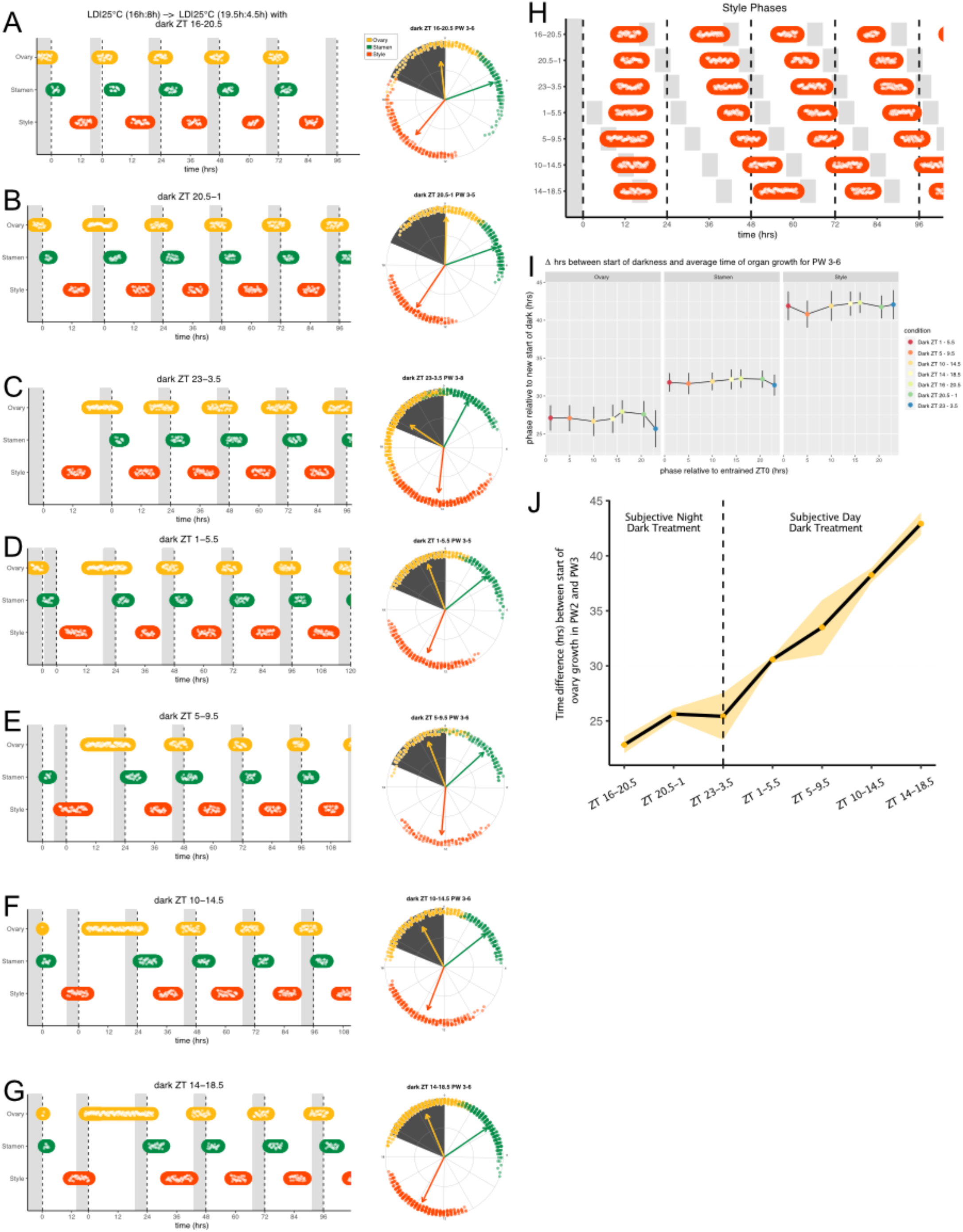
Phase of Organ Growth Resets Rapidly to New Entrainment Conditions. Plants were transitioned from LD|25°C (16h:8h) to 19.5h light:4.5 h dark at various times of the subjective day or night. New times of dark onset are indicated with grey boxes and labeled relative to the original entrainment condition. **(a-g)** Left plots are as described for Supplemental Fig 1. On the right plots, the timing of active organ growth for pseudowhorls 3 and later are superimposed on a 24-hour circular clock. Dots represent individual observations of growth, arrow angles represent average phases of growth, and arrow lengths represents precision of timing. (n=3-4 capitula for each condition). **(h)** Timing of active style growth in LD|25°C with the new short night at the indicated times (relative to the original entrainment condition; n = 3-4 capitula per condition). **(i)** Average times of growth for ovaries, stamens, and styles in pseudowhorls (PW) 3-6 for all indicated conditions. x-axis indicates phase of start-of-darkness for each condition relative to the original entrainment conditions. y-axis indicates phase of average organ growth relative to the start of the 4.5 hr dark treatment in the new entrainment conditions. Error bars indicate standard deviation. **(j)** Average difference in hours from the start of pseudowhorl 2 ovary growth and the start of pseudowhorl 3 ovary growth (y-axis) for each dark treatment condition (x-axis). Yellow dots represent mean differences, and the yellow ribbon represents standard deviation. Dotted line separates dark treatments given during the subjective night and the subjective day.

**Supplemental Figure 6.**
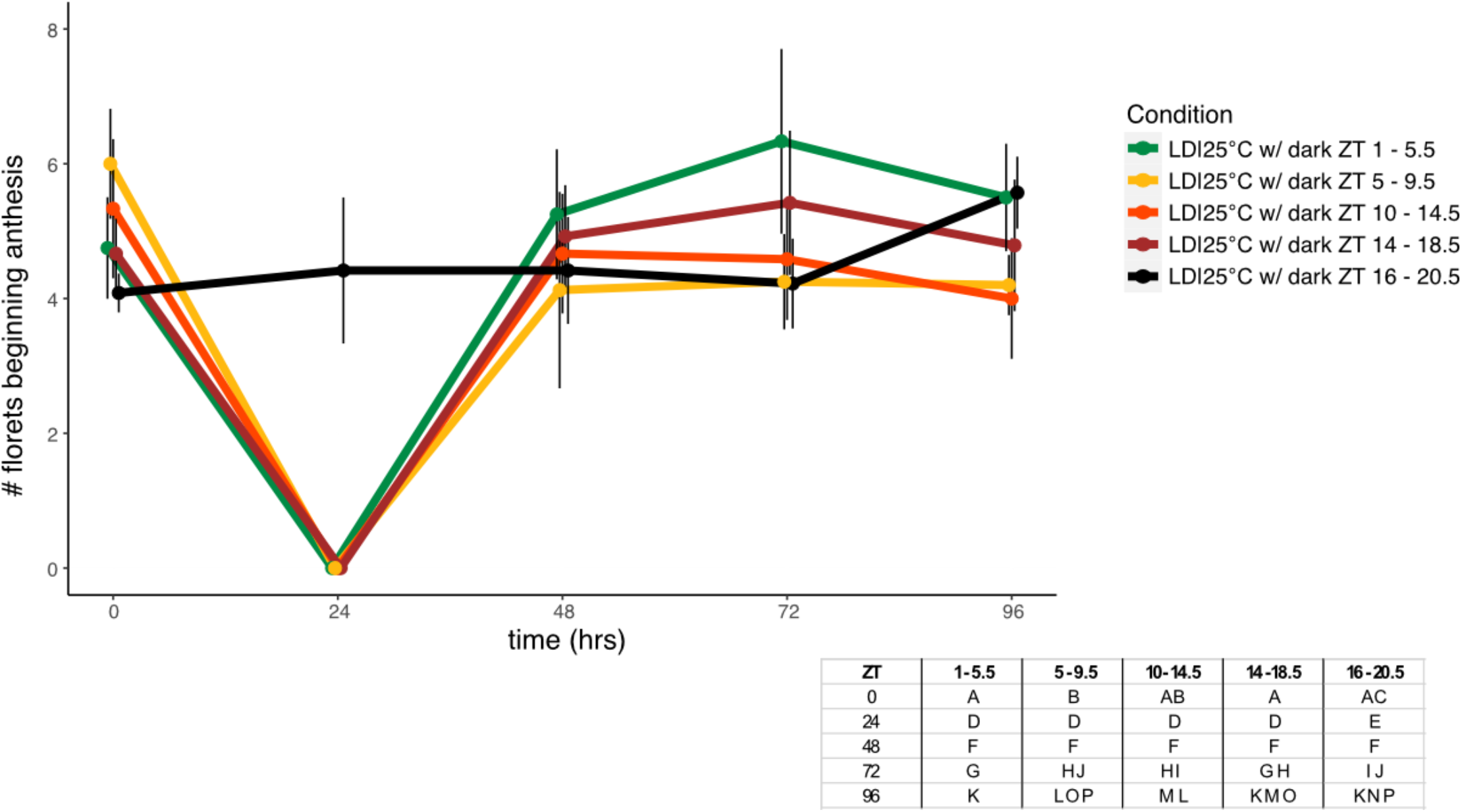
Number of Florets in a Pseudowhorl Not Altered After Delay in Anthesis. Average number of florets beginning anthesis along a parastichy and within a pseudowhorl for the indicated conditions (n = 12-19 parastichies per condition). At ZT 24, only condition LD|25°C with dark during ZT 16-20.5 had a new pseudowhorl undergoing anthesis. Error bars represent standard deviation. Different letters in table indicate statistically significant different groups within a time point, one-way ANOVA with post-hoc Tukey HSD test (*P* < 0.001).

**Supplemental Figure 7.**
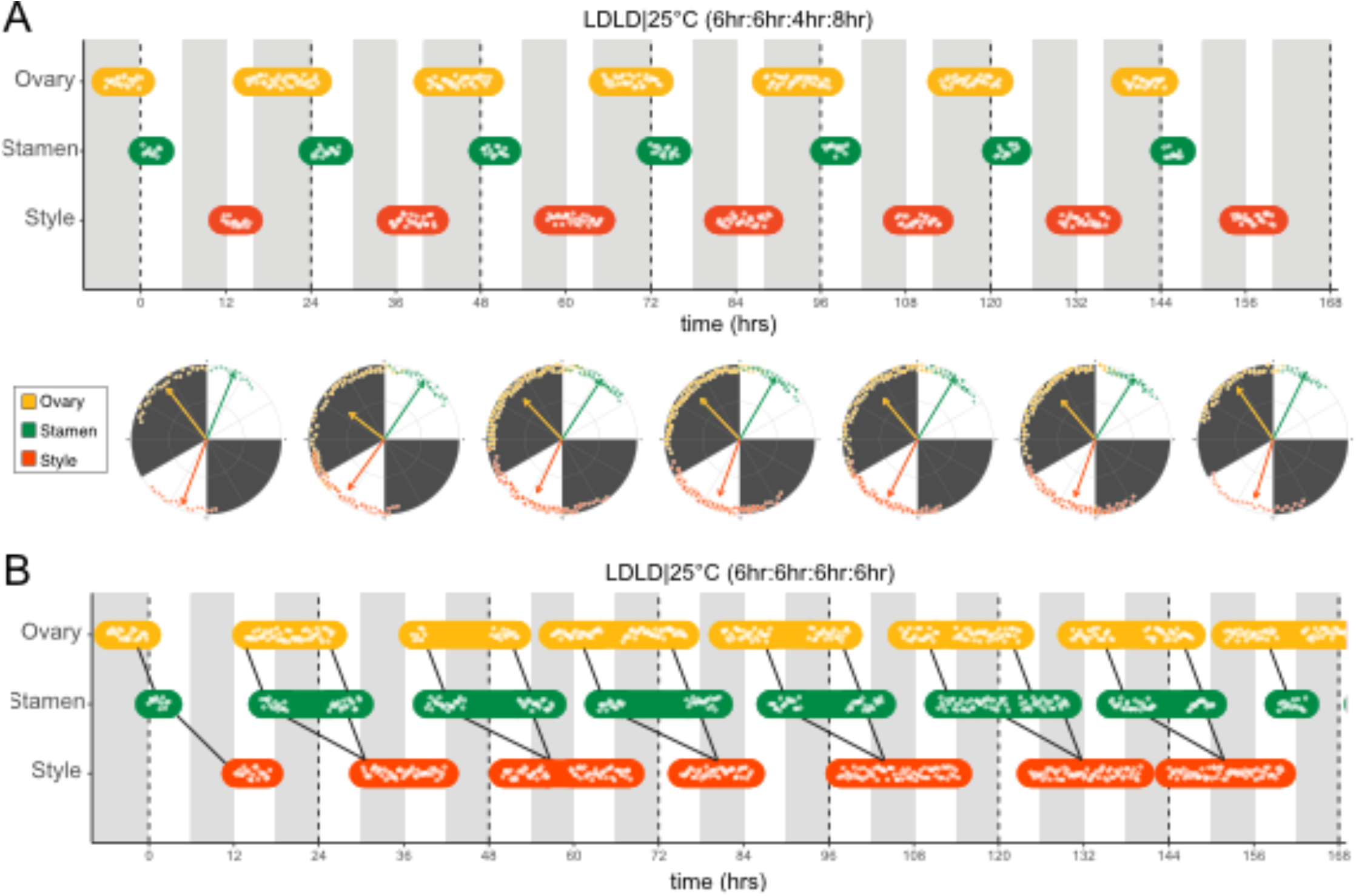
Frequency Demultiplication in Anthesis Requires a Dominant Night. Timing of ovary, stamen, and style growth for florets on a capitulum with two cycles of light and dark each day. **(a)** When one dark period is longer than the other (LDLD|25°C (6h L:6h D:4h L:8h D)), floret development only responds to the longer dark period. Individual polar plots are shown for each successive pseudowhorl. **(b)** When the two dark periods are of equal length (LDLD|25°C (6h L:6h D:6h L:6h D)), development is uncoordinated, with two distinct phases of growth within each pseudowhorl (n = 3 capitula per condition). See text for further description. Black lines link together floret organs within one pseudowhorl. Plots are otherwise as described for Supplemental Fig. 1.

**Supplemental Figure 8.**
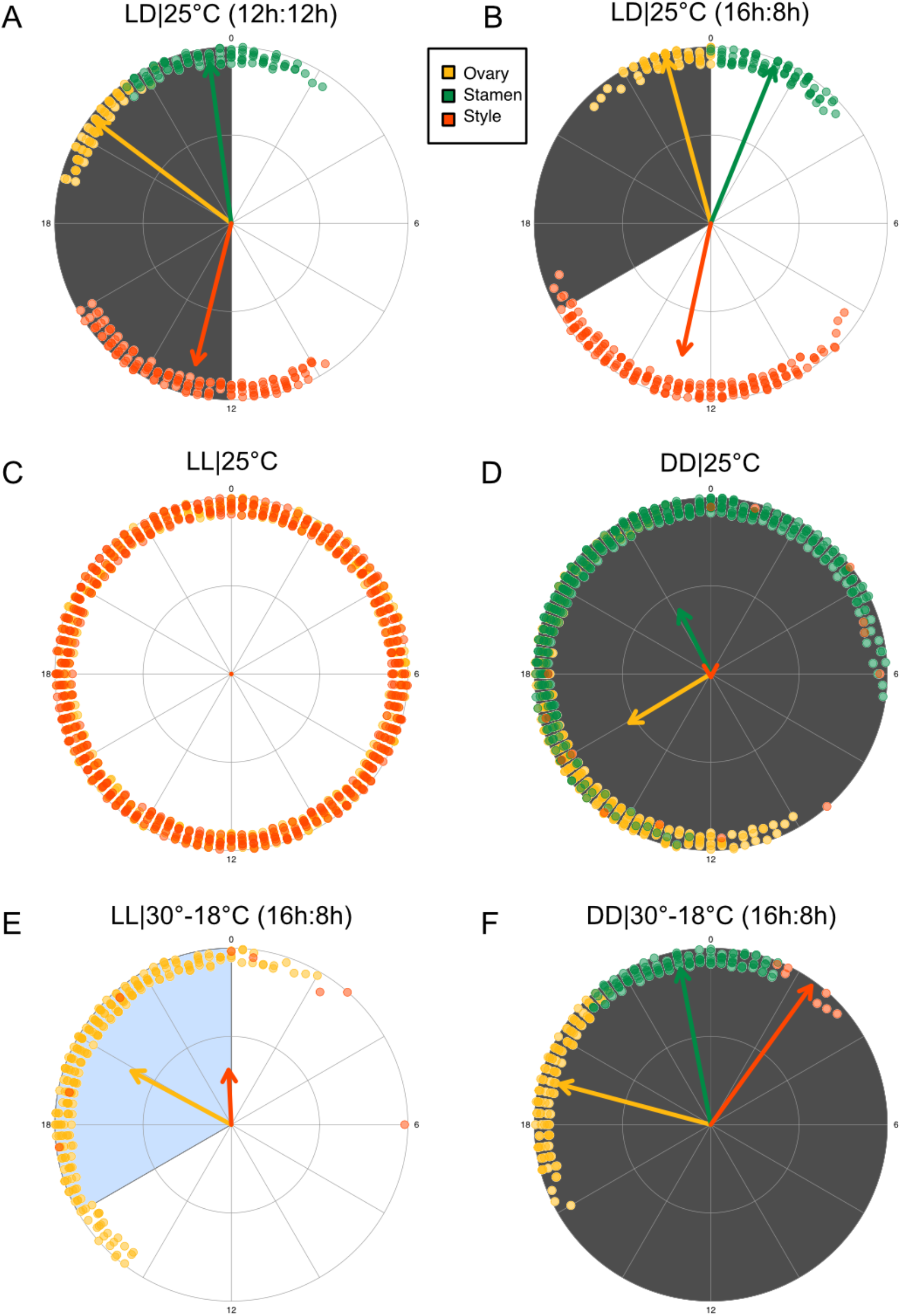
Relative Phases of Floret Development in Different Environmental Conditions. Phases after 48 hours in new entrainment conditions are depicted for organ growth in pseudowhorls 3 – 6 for conditions **(a)** LD|25°C (12h:12h), **(b)** LD|25°C (16h:8h), **(c)** LL|25°C, **(d)** DD|25°C, **(e)** LL|30°-18°C (16h:8h), and **(f)** DD|30°-18°C (n = 3-4 capitula per condition). Polar plots are as described for Supplemental Fig. 5.

**Supplemental Figure 9.**
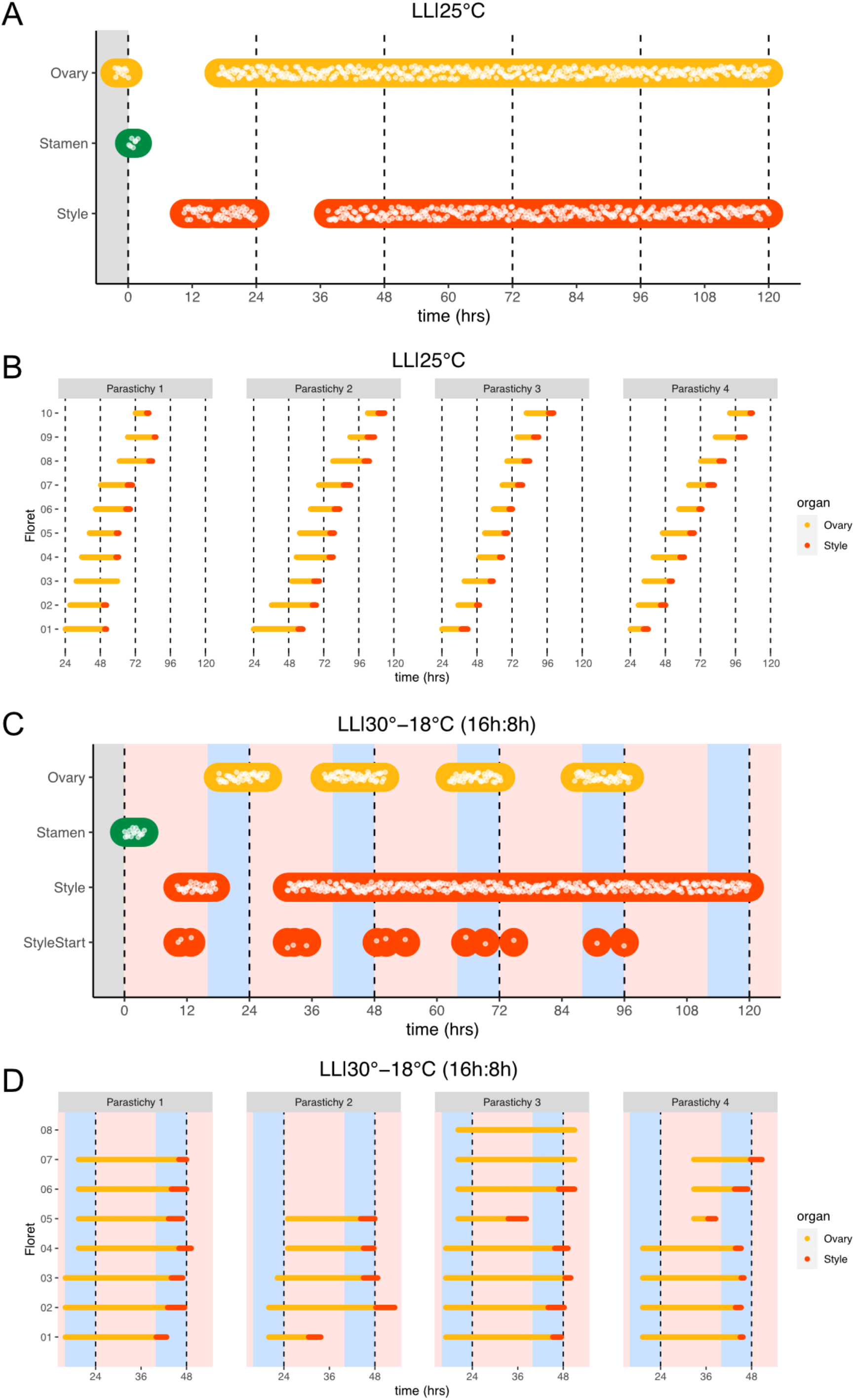
Sunflower Anthesis is Impaired in Constant Light. **(a, c)** Timing of ovary, stamen, and style growth for florets on a capitulum **(a)** LL|25°C (n=3 capitula) and **(c)** LL|30°-18°C (16h:8h) (n=3 capitula), representing growth for >5% of florets in a pseudowhorl (n=3). Plots are as described for Supplemental Fig. 1. **(b, d)** Timing of active growth of individual florets along a parastichy 24 hours after transfer to **(b)** LL|25°C for 10 florets per parastichy, and **(d)** LL|30°-18°C (16h:8h) for all florets along a parastichy within pseudowhorl 2 (n = 4 parastichies). Different temperatures are indicated by pink (30°C), white (25°C), and blue (18°C) backgrounds.

**Supplemental Figure 10.**
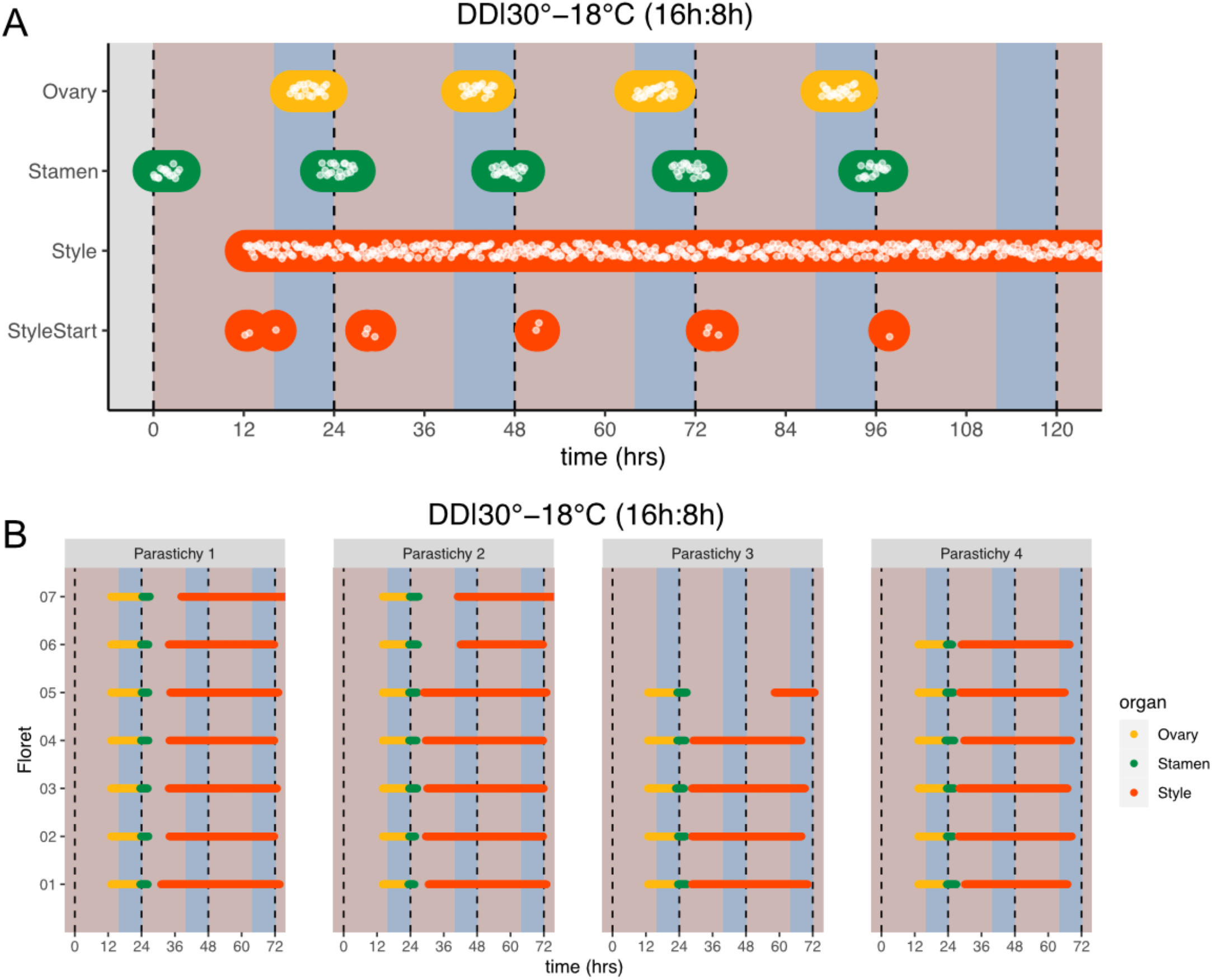
Thermocycles in Constant Dark Strengthen Coordination of Rhythms of Floret Development. **(a)** Timing of ovary, stamen, style, and start-of-style growth for florets on a capitulum in DD|30°-18°C (16h:8h) (n=3 capitula). **(b)** Timing of active growth for all florets along a parastichy within pseudowhorl 2 after transfer to DD|30°-18°C (16h:8h) (n = 4 parastichies). Plots are as described for Supplemental Fig 1.

**Supplemental Table 1.**
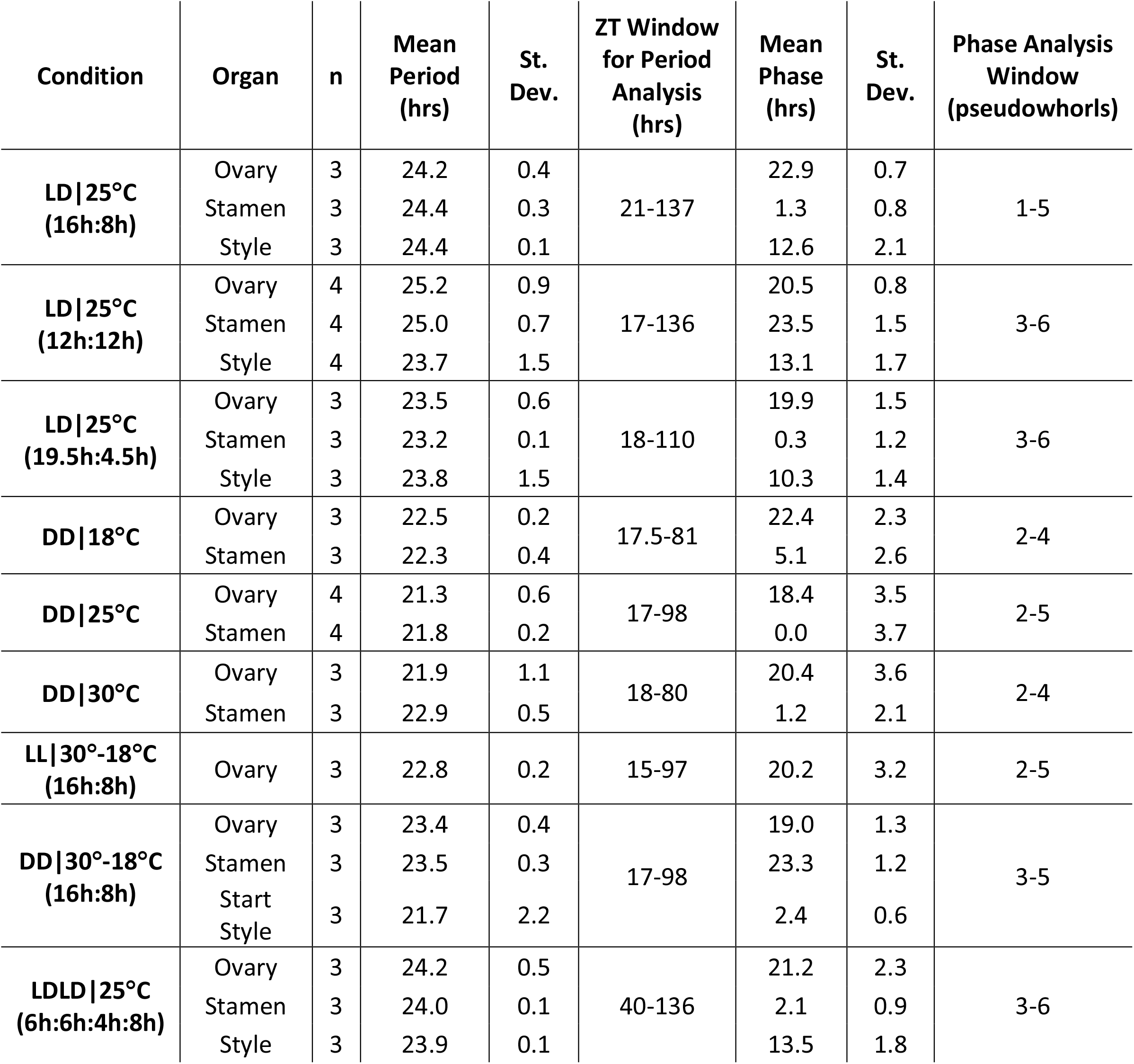
Circadian Parameters for Anthesis in Different Environmental Conditions.

**Supplemental Table 2.** Time Between Initiation of PW2 and PW3.

| **Period of Darkness (ZT)** | **Mean Time**<br>**Between PW2**<br>**and PW3 (hrs)** | **St. Dev.** |
| --- | --- | --- |
| **16-20.5** | 22.8 | 0.76 |
| **20.5-1** | 25.6 | 0.53 |
| **23-3.5** | 25.4 | 2.13 |
| **1-5.5** | 30.6 | 0.29 |
| **5-9.5** | 33.5 | 2.47 |
| **10-14.5** | 38.3 | 0.66 |
| **14-18.5** | 42.9 | 1.00 |

## Notes

### Competing Interest Statement

The authors have declared no competing interest.

## References

Agostinelli, C., Lund U. 2022. R package “circular”: Circular Statistics (version 0.4-94).

Amorim MD, Costa D da S, Krahl DRP, Fischer E, Rech AR. 2021. Gongylolepis martiana, an Asteraceae pollinated by bats in the Amazon. Plant Biol 23:728–734. doi:10.1111/plb.13283

Atamian HS, Creux NM, Brown EA, Garner AG, Blackman BK, Harmer SL. 2016. Circadian regulation of sunflower heliotropism, floral orientation, and pollinator visits. Science 353:587–590. doi:10.1126/science.aaf9793

Baroncelli S, Lercari B, Cecconi F, Pugliesi C. 1990. Light control of elongation of filament in sunflower (Helianthus annuus L.). Photochem Photobiol 52:229–231.

Bloch G, Bar-Shai N, Cytter Y, Green R. 2017. Time is honey: circadian clocks of bees and flowers and how their interactions may influence ecological communities. Philos Trans R Soc B Biol Sci 372:20160256. doi:10.1098/rstb.2016.0256

Brady D, Saviane A, Cappellozza S, Sandrelli F. 2021. The Circadian Clock in Lepidoptera. Front Physiol 12:776826. doi:10.3389/fphys.2021.776826

Budumajji U, Raju AJS. 2018. Pollination ecology of Bidens pilosa L. (Asteraceae). Taiwania 63:89–100.

Bünsow R. 1953. Endogene Tagesrhythmik und Photoperiodismus bei Kalanchoë blössfedliana. Planta 42:220–252.

Chandler JW. 2011. The Hormonal Regulation of Flower Development. J Plant Growth Regul 30:242–254. doi:10.1007/s00344-010-9180-x

Creux NM, Brown EA, Garner AG, Saeed S, Scher CL, Holalu SV, Yang D, Maloof JN, Blackman BK, Harmer SL. 2021. Flower orientation influences floral temperature, pollinator visits and plant fitness. New Phytol 232:868–879. doi:10.1111/nph.17627

DeGrandi-Hoffman G, Watkins JC. 2000. The foraging activity of honey bees Apis mellifera and non-Apis bees on hybrid sunflowers (Helianthus annuus) and its influence on cross-pollination and seed set. J Apic Res 39:37–45. doi:10.1080/00218839.2000.11101019

Dodd AN, Salathia N, Hall A, Kévei E, Tóth R, Nagy F, Hibberd JM, Millar AJ, Webb A a R. 2005. Plant circadian clocks increase photosynthesis, growth, survival, and competitive advantage. Science 309:630–633. doi:10.1126/science.1115581

Emerson KJ, Bradshaw WE, Holzapfel CM. 2008. Concordance of the circadian clock with the environment is necessary to maximize fitness in natural populations. Evolution 62:979–83. doi:10.1111/j.1558-5646.2008.00324.x

Funk VA, Susanna A, Stuessy TF, Robinson H. 2009. Classification of Compositae In: Funk VA, Susanna A, Stuessy TF, Bayer RJ, editors. Systematics, Evolution and Biogeography of the Compositae. Vienna: IAPT. pp. 171–189.

Greenham K, McClung CR. 2015. Integrating circadian dynamics with physiological processes in plants. Nat Rev Genet 16:598–610. doi:10.1038/nrg3976

Hipólito J, Roque N, Galetto L, Viana BF, Kevan PG. 2013. The pollination biology of Pseudostifftia kingii H.Rob. (Asteraceae), a rare endemic Brazilian species with uniflorous capitula. Braz J Bot 36:247–254. doi:10.1007/s40415-013-0023-4

Ikegami K, Yoshimura T. 2012. Circadian clocks and the measurement of daylength in seasonal reproduction. Mol Cell Endocrinol 349:76–81. doi:10.1016/j.mce.2011.06.040

Imaizumi T, Kay SA. 2006. Photoperiodic control of flowering: not only by coincidence. Trends Plant Sci 11:550–558. doi:10.1016/j.tplants.2006.09.004

Inoue K, Araki T, Endo M. 2018. Circadian clock during plant development. J Plant Res 131:59–66. doi:10.1007/s10265-017-0991-8

Izawa T, Oikawa T, Sugiyama N, Tanisaka T, Yano M, Shimamoto K. 2002. Phytochrome mediates the external light signal to repress FT orthologs in photoperiodic flowering of rice. Genes Dev 16:2006–2020. doi:10.1101/gad.999202

Jeffrey C. 2009. Evolution of Compositae Flowers In: Funk VA, Susanna A, Stuessy TF, Bayer RJ, editors. Systematics, Evolution and Biogeography of the Compositae. Vienna: IAPT. pp. 131–138.

Koritala BSC, Wager C, Waters JC, Pachucki R, Piccoli B, Feng Y, Scheinfeldt LB, Shende SM, Park S, Hozier JI, Lalakia P, Kumar D, Lee K. 2020. Habitat-Specific Clock Variation and Its Consequence on Reproductive Fitness. J Biol Rhythms 35:134–144. doi:10.1177/0748730419896486

Larrondo LF, Canessa P. 2018. The Clock Keeps on Ticking: Emerging Roles for Circadian Regulation in the Control of Fungal Physiology and Pathogenesis In: Rodrigues ML, editor. Fungal Physiology and Immunopathogenesis, Current Topics in Microbiology and Immunology. Cham: Springer International Publishing. pp. 121–156. doi:10.1007/82_2018_143

Lindström LI, Hernández LF. 2015. Developmental morphology and anatomy of the reproductive structures in sunflower (Helianthus annuus): a unified temporal scale. Botany 93:307–316. doi:10.1139/cjb-2014-0245

Lindström LI, Pellegrini CN, Hernández LF. 2007. Histological development of the sunflower fruit pericarp as affected by pre- and early post-anthesis canopy shading. Field Crops Res 103:229–238. doi:10.1016/j.fcr.2007.06.005

Lobello G, Fambrini M, Baraldi R, Lercari B, Pugliesi C. 2000. Hormonal influence on photocontrol of the protandry in the genus Helianthus. J Exp Bot. doi:10.1093/jxb/51.349.1403

Mandel JR, Dikow RB, Siniscalchi CM, Thapa R, Watson LE, Funk VA. 2019. A fully resolved backbone phylogeny reveals numerous dispersals and explosive diversifications throughout the history of Asteraceae. Proc Natl Acad Sci 116:14083–14088. doi:10.1073/pnas.1903871116

Marc J, Palmer JH. 1981. Photoperiodic sensitivity of inflorescence initiation and development in sunflower. Field Crops Res 4:155–164. doi:10.1016/0378-4290(81)90065-4

Marciniak, Przedniczek. 2019. Comprehensive Insight into Gibberellin- and Jasmonate-Mediated Stamen Development. Genes 10:811. doi:10.3390/genes10100811

McClung CR. 2019. The Plant Circadian Oscillator. Biol Basel 8. doi:10.3390/biology8010014

Michael TP, Mockler TC, Breton G, McEntee C, Byer A, Trout JD, Hazen SP, Shen R, Priest HD, Sullivan CM, Givan SA, Yanovsky M, Hong F, Kay SA, Chory J. 2008. Network Discovery Pipeline Elucidates Conserved Time-of-Day–Specific cis-Regulatory Modules. PLoS Genet 4:e14. doi:10.1371/journal.pgen.0040014

Michael TP, Salomé PA, Yu HJ, Spencer TR, Sharp EL, McPeek MA, Alonso JM, Ecker JR, McClung CR. 2003. Enhanced Fitness Conferred by Naturally Occurring Variation in the Circadian Clock. Science 302:1049–1053. doi:10.1126/science.1082971

Miller BH, Olson SL, Turek FW, Levine JE, Horton TH, Takahashi JS. 2004. Circadian Clock Mutation Disrupts Estrous Cyclicity and Maintenance of Pregnancy. Curr Biol 14:1367–1373. doi:10.1016/j.cub.2004.07.055

Mosier AE, Hurley JM. 2021. Circadian Interactomics: How Research Into Protein-Protein Interactions Beyond the Core Clock Has Influenced the Model of Circadian Timekeeping. J Biol Rhythms 36:315–328. doi:10.1177/07487304211014622

Muroya M, Oshima H, Kobayashi S, Miura A, Miyamura Y, Shiota H, Onai K, Ishiura M, Manabe K, Kutsuna S. 2021. Circadian Clock in Arabidopsis thaliana Determines Flower Opening Time Early in the Morning and Dominantly Closes Early in the Afternoon. Plant Cell Physiol 62:883–893. doi:10.1093/pcp/pcab048

Neff JL, Simpson BB. 1990. The Roles of Phenology and Reward Structure in the Pollination Biology of Wild Sunflower (Helianthus annuus L., Asteraceae). Isr J Bot 39:197–216.

Ouyang Y, Andersson CR, Kondo T, Golden SS, Johnson CH. 1998. Resonating circadian clocks enhance fitness in cyanobacteria. Proc Natl Acad Sci U A 95:8660–4. doi:10.1073/pnas.95.15.8660

Palmer JH, Steer BT. 1985. The generative area as the site of floret initiation in the sunflower capitulum and its integration to predict floret number. Field Crops Res 11:1–12. doi:10.1016/0378-4290(85)90088-7

Patterson B. 2009. Systematics, Evolution, and Biogeography of Compositae, Madroño. doi:10.3120/0024-9637-56.3.209

Pittendrigh CS, Minis DH. 1964. The Entrainment of Circadian Oscillations by Light and Their Role as Photoperiodic Clocks. Am Nat 98:261–294. doi:10.1086/282327

Putt ED. 1940. Observations on Morphological Characters and Flowering Processes in the Sunflower (Helianthus annuus L.). Sci Agric 21:167–179.

Rensing L, Ruoff P. 2002. Temperature effect on entrainment, phase shifting, and amplitude of circadian clocks and its molecular bases. Chronobiol Int 19:807–864. doi:10.1081/CBI-120014569

Richmond DL, Oates AC. 2012. The segmentation clock: inherited trait or universal design principle? Curr Opin Genet Dev 22:600–606. doi:10.1016/j.gde.2012.10.003

Roenneberg T, Dragovic Z, Merrow M. 2005. Demasking biological oscillators: properties and principles of entrainment exemplified by the Neurospora circadian clock. Proc Natl Acad Sci U S A 102:7742–7. doi:10.1073/pnas.0501884102

Ruan W, Yuan X, Eltzschig HK. 2021. Circadian rhythm as a therapeutic target. Nat Rev Drug Discov 20:287–307. doi:10.1038/s41573-020-00109-w

Sargent ML, Briggs WR, Woodward DO. 1966. Circadian Nature of a Rhythm Expressed by an Invertaseless Strain of Neurospora crassa. Plant Physiol 41:1343–1349. doi:10.1104/pp.41.8.1343

Schneider CA, Rasband WS, Eliceiri KW. 2012. NIH Image to ImageJ: 25 years of image analysis. Nat Methods 9:671–5.

Seiler GJ. 2015. Anatomy and Morphology of Sunflower In: Schneiter AA, editor. Agronomy Monographs. Madison, WI, USA: American Society of Agronomy, Crop Science Society of America, Soil Science Society of America. pp. 67–111. doi:10.2134/agronmonogr35.c3

Spoelstra K, Wikelski M, Daan S, Loudon ASI, Hau M. 2016. Natural selection against a circadian clock gene mutation in mice. Proc Natl Acad Sci 113:686–691. doi:10.1073/pnas.1516442113

Valentin-Silva A, Godinho M, Cruz K, Lelis S, Vieira M. 2016. Three psychophilous Asteraceae species with distinct reproductive mechanisms in southeastern Brazil. N Z J Bot 54:498–510. doi:10.1080/0028825X.2016.1236735

Visscher PK, Seeley TD. 1982. Foraging Strategy of Honeybee Colonies in a Temperate Deciduous Forest. Ecology 63:1790. doi:10.2307/1940121

Wenden B, Kozma-Bognár L, Edwards KD, Hall AJW, Locke JCW, Millar AJ. 2011. Light inputs shape the Arabidopsis circadian system. Plant J 66:480–491. doi:10.1111/j.1365-313X.2011.04505.x

Wickham H, Averick M, Bryan J, Chang W, McGowan L, François R, Grolemund G, Hayes A, Henry L, Hester J, Kuhn M, Pedersen T, Miller E, Bache S, Müller K, Ooms J, Robinson D, Seidel D, Spinu V, Takahashi K, Vaughan D, Wilke C, Woo K, Yutani H. 2019. Welcome to the Tidyverse. J Open Source Softw 4:1686. doi:10.21105/joss.01686

Xuan W, De Gernier H, Beeckman T. 2020. The dynamic nature and regulation of the root clock. Development 147:dev181446. doi:10.1242/dev.181446

Yanovsky MJ, Kay SA. 2003. Living by the calendar: how plants know when to flower. Nat Rev Mol Cell Biol 4:265–276. doi:10.1038/nrm1077

Yon F, Joo Y, Cortés Llorca L, Rothe E, Baldwin IT, Kim S-G. 2016. Silencing Nicotiana attenuata LHY and ZTL alters circadian rhythms in flowers. New Phytol 209:1058–1066. doi:10.1111/nph.13681

Yon F, Kessler D, Joo Y, Cortés Llorca L, Kim SG, Baldwin IT. 2017. Fitness consequences of altering floral circadian oscillations for Nicotiana attenuata. J Integr Plant Biol 59:180–189. doi:10.1111/jipb.12511

Zhang T, Cieslak M, Owens A, Wang F, Broholm SK, Teeri TH, Elomaa P, Prusinkiewicz P. 2021. Phyllotactic patterning of gerbera flower heads. Proc Natl Acad Sci 118:e2016304118. doi:10.1073/pnas.2016304118

Zielinski T, Moore AM, Troup E, Halliday KJ, Millar AJ. 2014. Strengths and limitations of period estimation methods for circadian data. PLoS ONE 9:e96462. doi:10.1371/journal.pone.0096462 PONE-D-13-43174 [pii]

